# Odor Annoyance, Sensory Irritation or Relaxation: Acute Effects of Real Pinewood Emissions in Indoor Air Scenarios

**DOI:** 10.64898/2026.07.03.736270

**Authors:** Christine Ida Hucke, Viviane Gallus, Katja Butter, Julian Elias Reiser, Martin Ohlmeyer, Christoph van Thriel

## Abstract

Wood is commonly used in the building sector, emitting volatile organic compounds (VOCs) contributing to indoor air quality. These VOC profiles can have a pleasant smell and positive effects e.g., induce relaxation. Contrarily, VOCs can have adverse health effects in higher concentrations. Therefore, some VOCs are regulated by guide values (GV). Potentially positive and negative effects of pinewood emissions, ranging from 0.2 mg/m^3^ (German GV I for bicyclic terpenes) to 2.0 mg/m^3^ (GV II) were investigated in an experimental 2 h exposure study using a within-subject design. Thirty-two healthy participants rated the perception, pleasantness, symptoms of irritation, and indicators of well-being. During a demanding working memory task (n-back) and a resting period, heart rate (HR) and HR variability (HRV) changes were measured. Before and after each session physiological markers of sensory irritation were assessed. Ratings indicated that the exposure to GV I and GV II were not perceived as more intense or pleasant. Mostly concentration-independent effects were revealed, indicating that inter-individual factors influenced the ratings rather than the VOCs. The pinewood odors during the n-back task did not cause distraction nor did it facilitate performance as previously suggested. HR/V changes indicated that pinewood odors during and after the n-back tasks did not induce relaxation. Only symptoms of nasal irritation showed some weak concentration-dependency, not supported by physiological markers or comparable ratings of sensory irritation. In conclusion, the fact that no distinct odor is detected suggests that interfering factors potentially prevent the regulation of odors at relevant indoor air concentrations.

**Highlights:**

- Successful exposure to realistic indoor pinewood VOCs
- No effect on ratings of indoor air quality, performance or sensory irritation
- No induction of relaxation during or after stressful tasks
- Short-term exposure to concentrations between GV I and II are non-effective

## 1 Introduction

Wood is a popular construction material also used for indoor interior design due to its sustainability and biophilic design. Softwoods (e.g., pinewood) are among the most popular wood types for construction or furniture in Germany [1] and other European countries such as Finland [2], [3] as well as Northern America [4]. As an organic material, wood emits volatile organic compounds (VOCs). The highest proportion of the pinewood emission profile are terpenes, with α-pinene, 3-carene, β-pinene, and limonene making up the highest proportion of these terpenes [5], [6], [7]. Therefore, these VOCs are relevant for the evaluation of indoor air quality.

The German Committee for Indoor Air Quality Guidelines (AIR) established indoor air guide values (GV) to avoid adverse health effects indoors. The GV I value (precautionary guide value) is the concentration that should be undershot long term and if doing so, no negative health effects are to be expected. The GV II value (hazard guide value) is an effect-related value that is derived from toxicological studies. Here, the no-observed-adverse-effect level (NOAEL) or lowest-observed-adverse-effect level (LOAEL) are used and by applying various assessments factors (e.g., inter-and intra-species differences) an airborne concentration serving as GV I or II is derived. Thus, GV II is a health-based limit value and if this concentration is exceeded, immediate actions are required to reduce the VOC concentration to avoid any adverse health risk for the room users. The range between GV I and GV II is not considered to be safe and counteractive measures (e.g., increased ventilation or cleaning) might be required to diminish the undesirable contamination. Often, GV I is derived from the GV II using an additional factor (e.g., of 10) [8], [9]. GV values for bicyclic terpenes (i.e., α-pinene, 3-carene, and β-pinene) were derived from a LOAEL of 450 mg/m^3^ that elicited sensory irritation in human volunteers [10]. Thus, the GVs range from 0.2 mg/m^3^ (GV I) to 2 mg/m^3^ (GV II). Recently, specific values of 0.5 mg/m^3^ (GV I) and 1 mg/m^3^ (GV II) for α-pinene have been established by the German AIR [11], using an inhalation study in rodents as point of departure while using a different approach when deriving the GV I and II. In contrast to the previous justification of the bicyclic terpenes document the GV I was derived from the NOAEL of a 14-week inhalation study in mice and rats, while GV II was derived from the LOAEL of this NTP study [12]. While these values are occasionally surpassed, e.g., during active building phases or remodeling [13], [14], realistic indoor air values usually fall within the range of GV I and II and are, for the most part, even below the GV I value [14], [15], [16], [17], [18], [19], [20]. However, even experimental studies exceeding the GV II value during short-term exposures did not report any signs of sensory irritation in concentrations up to 9.5 mg/m^3^ [21]. Further, no risk to develop asthma or aggravate existing symptoms was posed due to exposure at higher concentrations as shown in an experimental mice study as well as over time at lower concentrations as supported by a large-scale cohort study [22].

The above-mentioned concerns are based on sensory irritation that can be evoked by VOCs binding to receptors of the trigeminal nerve innervating the mucous membranes of the nose, the upper respiratory tract and the conjunctiva. This cranial nerve (CN V) is responsible for irritating perceptions such as burning, stinging but also coolness of e.g. menthol, depending on the receptor they bind to [23], [24]. Further, reflexes such as sneezing or eye blinks are triggered, preventing the body from potential harm [25] as prolonged exposure might result in neurogenic inflammation and subsequent tissue damage [26]. Additionally, VOCs can bind to olfactory receptor neurons in the nose and evoke olfactory perceptions [27]. These perceptions can range from pleasant to unpleasant [24]. Thus, even if a VOC is not considered to be irritating, a malodor might still evoke unpleasant olfactory perceptions [28]. Some VOCs have been reported to evoke undesirable reactions such as nausea or headache based on olfactory properties, at least in some individuals. Recently, attempts have been on the forefront to prevent odor annoyance-related (hygienic) negative effects by using odor guide values (OGV) [29], [30]. Another assumption is that especially malodors and unpleasant perceptions might impair work performance or increase mental load when performing cognitive tasks [31]. Such behavioral or neurophysiological effects of annoying odors have been proposed as a proxy for assessing adverse effects of odors at least in the working environment [26]. Research investigating potential direct odor-related negative effects in concentrations that are below the irritation threshold and thereby purely link to odor annoyance is lacking, though. Especially in the context of wood emissions, these aspects have not been systematically studied systematically.

In contrast to the potentially negative effects of wood emissions, wood and its odor have previously been linked to positive effects such as pleasant perceptions, relaxation and stress reduction or mood improvement [17], [32], [33], [34], [35], [36], [37], [38], [39], [40]. However, the stimulation setups used in these studies are partly artificial, bearing the question of how these effects relate to real-life indoor air scenarios. In a recent study, Jyske et al., [41] could show that olfactory perceptions of pinewood in rooms lacking visual wood features improved restoration outcome. However, visual and multisensory impressions had similar or additive effects, respectively. Furthermore, Muilu-Mäkelä et al., [17] showed that wooden offices with measurable terpene VOCs reduced anxiety levels while other parameters remained unchanged.

Clearly, there are contradictory reports concerning consequences of indoor wood VOCs [42]. The aim of the current study is to investigate the effects of real wood emissions in the concentrations of GV I and GV II to explore (a) potential positive effects of wood emissions and (b) potential annoyance or even sensory irritation effects evoked by the emissions. Multi-level full body exposure studies offer a standardized framework to explore the effects of exposure to specific VOCs in a controlled environment on the levels of perception, behavior and physiology while simulating an office-like workday situation (e.g., Kleinbeck et al., [43]).

Positive effects are the pleasant perceptions of the wood smell and the effects of relaxation, reflected in either subjective ratings or objective physiological parameters. Objectively, this is measured by investigating heart rate (HR) and heart rate variability (HRV) changes during a challenging working memory task and a subsequent resting period during the exposure with wood VOCs (or filtered air). As the body is regulated by the parasympathetic (normal function under relaxation) and sympathetic (activating during stress) nervous system [44], cardiac measures are helpful tools to derive information about processes of the nervous system. HRV parameters can be investigated in the time (e.g., standard deviation of all filtered heart beat (NN) intervals (SDNN) or square root of the mean of the sum of the squares of differences between adjacent NN intervals (RMSSD)) or frequency domain, i.e., low frequency (LF), high frequency (HF) or the ratio (LF/HF) indicating the dominance of either autonomous system [44], [45]. Following the Stress Reduction Theory (SRT) [46], nature and natural elements support recuperation from stress. For instance, exposure to wood scents or individual components such as α-pinene has been shown to not only improve mood but also HR and HRV parameters, indicating an induction of relaxation [34], [39], [47]. Thus, in the current study we expected that the wood odor might help to restore HR and HRV parameters to a more relaxed physiological state after stress induction by means of a working memory task. That is, HR and LF/HF decrease whereas HF, SDNN as well as RMSSD increase.

Negative effects might consolidate as annoyance due to the perceptions of smells or early signs of sensory irritation, which, again, were assessed through ratings of intensity of irritating perception and ratings of symptoms related to irritation. Physiological parameters that are used include eye redness, changes to the tear film stability, nasal concentrations of the neuropeptide Substance P [48] or the alarmin High-Mobility-Group-Protein B1 (HMGB1) [49], indicating the crosstalk of the peripheral nervous system and the immune system through neurogenic reflexes. An increase in fractionated exhaled nitric oxide (FeNO) is an early sign of upper airway inflammation [50]. Moreover, HR/HRV parameters might indicate induction of physiological stress due to exposure to GV I or GV II concentrations by showing a downregulation of the parasympathetic and upregulation of the sympathetic nervous system as reflected by an increase in HR and LF/HF and decrease in HF, SDNN and RMSSD.

Lastly, performance changes during a working memory task might indicate either negative or positive consequences of the emissions following the SRT as mediated by physiological relaxation [46] or direct effect on the attentional system as postulated by the Attention Restoration Theory (ART) [51]. This effect would manifest in either a performance boost or degradation as indicated by a significantly more or less accurate performance during the working memory task during the exposure compared to the control condition.

Using this multidimensional approach of endpoints, we investigate if the concentrations in the range of GV I – GV II for bicyclic terpenes are to be considered relevant for either positive (e.g., relaxation) or negative (e.g., annoyance or irritation) consequences.

## 2 Methods

The protocol was approved (approval #247) by the local ethics committee of the Leibniz Research Centre for Working Environment and Human Factors (IfADo). Participants gave written informed consent before the experiment started.

### 2.1 Participants

In total 33 participants were invited to take part in the experiment. One participant did not participate in all three conditions and was thus excluded from the analysis. Each test and analysis underwent an individual quality check (e.g., analytical detection level, participants’ incorrect response patterns during the n-back tasks, signal transmission issues during the HR recordings). In case further participants were excluded from specific analyses this is noted in the respective results sections. Table 1 indicates the demographic data and sample description including 32 participants.

**Table 1.** Demographic descriptions of the participants and questionnaire indices.

| Demographic |  |  |
| --- | --- | --- |
| N – Participants (complete 3 sessions) / ECG | 32 / 28 |  |
| Sex | $f = 20, m = 12$ | |
| Age | $M = 25.38, SD = 3.64$ | |
| Sniffin' Sticks – Total (TDI)* | $M = 34.94, SD = 3.39$ | Objective olfactory performance |
| Threshold (T) | $M = 9.81, SD = 2.13$ | |
| Discrimination (D) | $M = 12.41, SD = 1.66$ | |
| Identification (I) | $M = 12.56, SD = 1.63$ | |
| Questionnaires |  |  |
| CSS1 „Chemical Sensitivity Scale“ [53]* | $M = 29.81, SD = 6.23$ | Subjective olfactory sensitivity |
| Chemical Odor Sensitivity Scale (COSS) [54]* | $M = 14.03, SD = 9.11$ | Subjective olfactory sensitivity |
| Cacosmia Sum Score [55] | $M = 17.22, SD = 5.58$ | Questionnaire olfactory effects |
| Trait - State Trait Anxiety Inventory (STAI) | $M = 38.16, SD = 8.26$ | Index for anxiousness |
| Chemical and General Environmental Sensitivity Questionnaire (CGES) [56] |  | Subjective environmental and chemical sensitivity |
| General | $M = 39.50, SD = 12.60$ | |
| Body | $M = 7.69, SD = 4.28$ | |
| Respiration | $M = 19.78, SD = 11.22$ | |
| Skin/Allergy | $M = 17.59, SD = 16.87$ | |
| Odor Awareness Scale [57] – total OAS | $M = 120.50, SD = 18.03$ | Subjective daily odor presence |
| Negative Odor Awareness | $M = 74.63, SD = 11.00$ | |
| Positive Odor Awareness | $M = 45.84, SD = 7.83$ | |
| NEO FFI [58] |  | Personality questionnaire |
| Neuroticism | $M = 1.70, SD = 0.64$ | |
| Extraversion | $M = 2.32, SD = 0.57$ | |
| Openness for experiences | $M = 2.81, SD = 0.56$ | |
| Agreeableness | $M = 2.55, SD = 0.54$ | |
| Conscientiousness | $M = 2.69, SD = 0.66$ | |

N = sample size, ECG = electrocardiogram
\* Sniffin' Sticks results indicate normal olfactory function compared to previous experiments (e.g., [59], [60], [61] and show normal values in the final sample description [53], [54], [62].

Before participation, potential participants were screened in a phone interview. Exclusion criteria were pregnancy, regular smoking or drug consumption, migraine, medical conditions of the cardiopulmonary system (e.g., asthma) or the central nervous system, certain allergies (e.g., acute hay fever), or certain hairstyles that prevent an adequate EEG recording (not relevant for the current analysis).

All participants completed a “training day” (see section 2.4.1) to familiarize themselves with the task and procedures during the experimental exposures and to assess their olfactory acuity and personality traits (see Table 1) that might affect assessment of acute symptoms and perception [52].

Overall, the sample had a slightly higher proportion of female participants. In general, all participants unimpaired olfaction, no signs of self-reported hypersensitivity towards odors, and normal personality traits with respect to standardized questionnaire [58].

### 2.2 Exposure

#### 2.2.1 Full Body Exposure Chamber (ExpoLab)

The exposure was conducted in the environmental chamber at the IfADo (ExpoLab) designed to perform controlled human exposure studies (Figure S 1). The ExpoLab is a 28 m^3^ room built of glass and stainless steel to prevent absorption and emission of VOCs. The lab can be entered through an airlock system, preventing a change in VOC concentration. The lab is equipped with a ventilation system (5-fold air change/hour, mean temperature: 23°C, humidity: 45.8 %) that is connected to an air conditioning unit in an adjacent room. Via a branched pipe system on the bottom of the ExpoLab this system supplies the room with conditioned air that can be enriched with desired VOCs in a predefined concentration. The ExpoLab is operated with slight negative pressure to minimize leakage into the surrounding laboratory and to direct the airflow to the roof of the room. Here, four sampling pipes which are connected to a GC-FID (PerkinElmer Clarus® 690 GC) in the adjacent control room ensure a quasi-continuous air monitoring of the VOC-enriched air. Every 2.5 minutes samples are drawn and analyzed, and the target concentration can be adjusted accordingly.

#### 2.2.2 Exposure Scenarios

Shredded pinewood chips (*Pinus sylvestris* L.) were used to create a natural VOC profile. The pinewood chips were produced in one batch by members of the Thünen Institute and frozen until used at the IfADo. To generate the VOCs for the exposures, 200 L oven bags were filled with 200 g of pinewood chips and carefully heated to 50°C for 1-1.5 h. Subsequently, the VOC-enriched air was filtered and evacuated to second bag of the same size. Samples of each of these bags were analyzed using GC-FID (PerkinElmer Clarus® 690 GC; calibrated to α-pinene and 3-carene) to ensure the desired concentration of α-pinene and 3-carene (between 5.750 and 11.500 mg/m^3^). This second bag with a known VOC concentration was then connected to the inlet airstream of the climate control ventilation system and diluted to create three exposure scenarios:

- GV I for bicyclic monoterpenes (0.2 mg/m^3^)
- GV II for bicyclic monoterpenes (2.0 mg/m^3^)
- Control exposure with filtered air

The GC-FID in the control room measured the respective concentration throughout the experiment. If necessary, the dosing from the sample bag could be adjusted to ensure the intended and stable concentration throughout the experiment. The combined α-pinene and 3-carene concentrations measured by online GC-FID during the experiment averaged 0.23 mg/m³ (SD = 0.004 mg/m³) in the GV I scenario and 2.27 mg/m³ (SD = 0.04 mg/m³) in the GV II scenario (Supplement 2). To establish this protocol, several test runs were conducted while taking air samples using Tenax TA®-tubes which were sent to the Thünen Institute and analyzed using gas-chromatography-mass spectrometry (Supplement 3A). During the experimental phase these measurements were repeated twice per condition to validate a stable concentration and VOC profile throughout the sessions. GC–MS analyses of sampled air confirmed that the air in the ExpoLab was predominantly composed of α-pinene and 3-carene, with only minor additional concentrations of typical pine-wood-related terpenes, including β-pinene and limonene (Supplement 3B).

Considering the recently published guideline values for α-pinene alone and the subsequent withdrawal of guideline values for bicyclic monoterpenes (defined as the sum of α-pinene, 3-carene, and β-pinene) [11], our results should also be interpreted in this context. The mean α-pinene concentrations during the study in the GV I and GV II scenarios were 0.121 mg/m³ (*SD* = 0.006 mg/m³) and 1.063 mg/m³ (*SD* = 0.057 mg/m³), respectively.

### 2.3 Measured endpoints

The following measurements were taken before and after or during each experimental exposure (See Table 2). To ensure a smooth execution, all measurements were explained by laboratory staff and practiced by the participants during the training day (See 2.4.1). In addition to the here-mentioned endpoints, samples of tear fluid as well as electroencephalographic (EEG) recordings were taken but will be published separately.

**Table 2.**
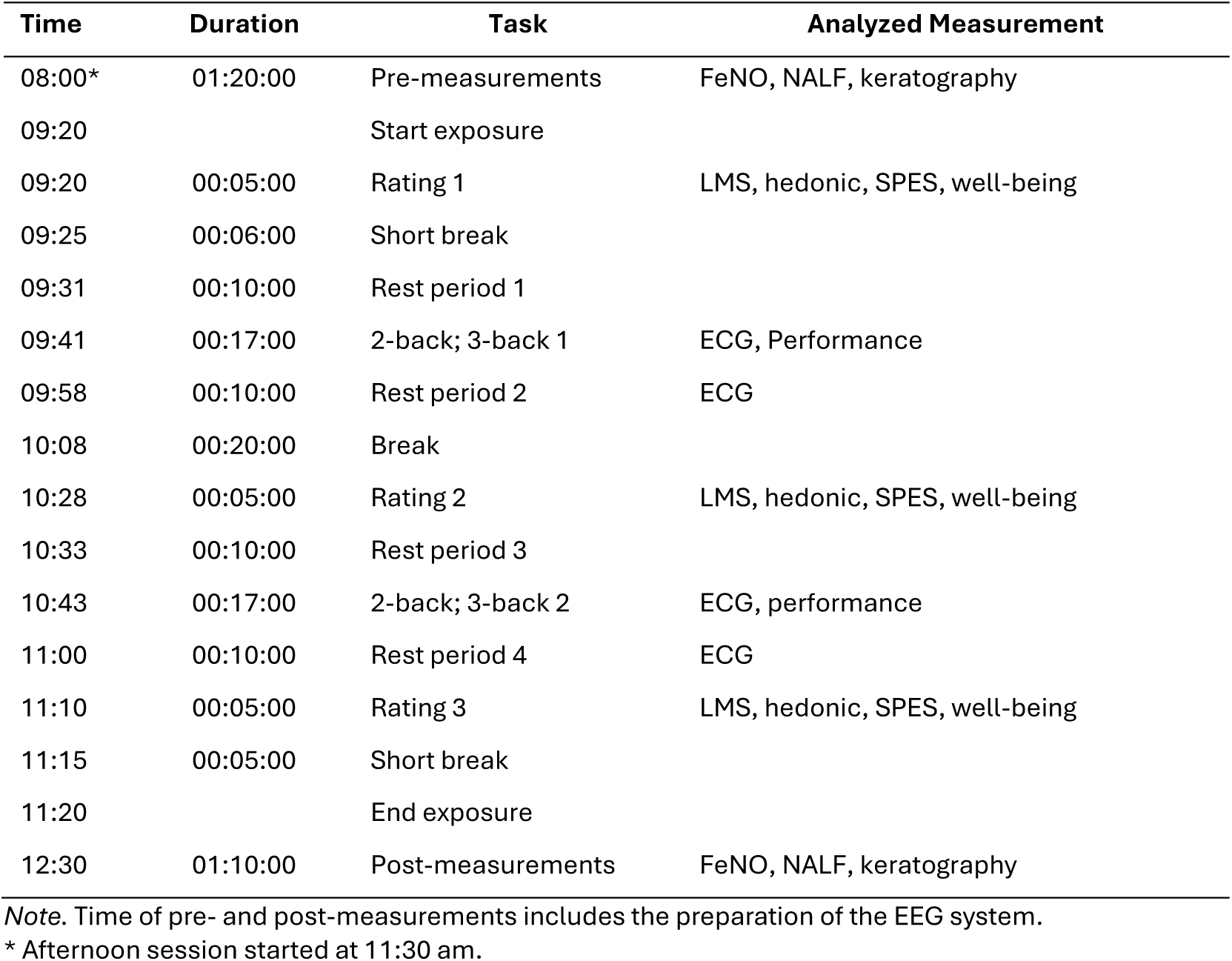
Schedule of a morning experimental exposure session.

| Time | Duration | Task | Analyzed Measurement |
| --- | --- | --- | --- |
| 08:00* | 01:20:00 | Pre-measurements | FeNO, NALF, keratography |
| 09:20 |  | Start exposure |  |
| 09:20 | 00:05:00 | Rating 1 | LMS, hedonic, SPES, well-being |
| 09:25 | 00:06:00 | Short break |  |
| 09:31 | 00:10:00 | Rest period 1 |  |
| 09:41 | 00:17:00 | 2-back; 3-back 1 | ECG, Performance |
| 09:58 | 00:10:00 | Rest period 2 | ECG |
| 10:08 | 00:20:00 | Break |  |
| 10:28 | 00:05:00 | Rating 2 | LMS, hedonic, SPES, well-being |
| 10:33 | 00:10:00 | Rest period 3 |  |
| 10:43 | 00:17:00 | 2-back; 3-back 2 | ECG, performance |
| 11:00 | 00:10:00 | Rest period 4 | ECG |
| 11:10 | 00:05:00 | Rating 3 | LMS, hedonic, SPES, well-being |
| 11:15 | 00:05:00 | Short break |  |
| 11:20 |  | End exposure |  |
| 12:30 | 01:10:00 | Post-measurements | FeNO, NALF, keratography |

*Note.* Time of pre- and post-measurements includes the preparation of the EEG system.
\* Afternoon session started at 11:30 am.

#### 2.3.1 Pre- and post-measurements (Physiological Irritation)

##### 2.3.1.1 Nasal Lavage

A nasal lavage was performed to collect potential proteins corresponding to neurogenic inflammation, i.e., Substance P and HMGB1 in the nasal lavage fluid (NALF). Using an electrical pipette, 5 ml of a sterile, isotonic (0.9 %) NaCl solution, warmed to 37°C, were inserted into each nostril while participants performed a velopharyngeal closure technique to seal the velum and pharyngeal walls, separating the oral and nasal cavities while breathing through the mouth. Participants were asked to keep the fluid inside the nostril for 10 seconds. Thereafter, the fluid was filtered for sediments and collected in an Eppendorf Tube and frozen at −80°C until analyzed. An Enzyme-linked Immunosorbent Assay (ELISA) was used to quantify Substance P (Cayman Kit 583751, Biomol GmbH, Hamburg, Germany) and HMGB1 (ST 51011, IBL International GmbH, Hamburg, Germany).

##### 2.3.1.2 Eye Irritation

A keratoscope (Keratograph 5M, OCULUS Optikergeräte GmbH, Wetzlar, Germany) was used to measure break-up time (BUT) of the precorneal tear film and eye redness. The BUT is an indicator for the stability of the tear film, while the eye redness is a software-based analysis of ocular bulbar redness indicating eye inflammation or dry eye disease [63]. BUT was measured at up to 144 corneal areas at both eyes. If there is no break-up of tear fluid, the measurement stopped automatically after 25 sec. As a result, times of first break-up (BUT1) are given (irrespective of the measured area). Furthermore, mean break-up is computed (BUTM), comprising all analyzed corneal areas. For eye redness analysis, pictures of the volunteer’s eyes were taken under standardized conditions. The method used by the R-Scan software is proprietary but shows a high linear relationship (R² = 0.99) to an analysis of the red channel of the eyes’ photos using a color extraction method described by Wolffsohn [64]. In this method, the relative intensity of red (R), green (G), and blue (B) color channels are determined in the bulbar area of the eyes. The Red-value in this method is determined by the ratio of the red-channel relative to the total color-channel activity (sum of R, G, and B channels). This procedure assures that the image brightness has no effect on the Red-Value (Downie et al. 2016). The R-Scan uses a clinical grading scale of 0-4 in 0.1 steps (referring to Jenvis grading scale). Nasal and temporal bulbar redness grades are provided automatically. The analyzed redness grade of each eye is the mean of temporal and nasal bulbar redness.

##### 2.3.1.3 FeNO

A NIOX VERO^®^ was used to measure the FeNO in the participants exhaled air. To that end, participants in- and exhaled through the mouthpiece of the NIOX VERO®. More specifically, participants were asked to slowly but constantly inhale until maximum capacity (or until a visual cue was given by the system) and then exhale for approximately 60 seconds until the NIOX VERO® gave a visual and audible cue to stop exhaling. The internal software indicated the FeNO in parts per billion (ppb). Applied at the mouth, NO-values correspond to the middle and lower airways. NO is produced in healthy airways. Normal values are lower than 25 ppb for adults [65], however, high values (higher than 50 ppb) indicate airway eosinophilia [65].

#### 2.3.2 During exposure

##### 2.3.2.1 Labeled Magnitude Scale

Throughout each exposure session, participants rated their subjective perceptions on the ‘Labeled Magnitude Scale’ (LMS) [66]. The LMS is a quasi-logarithmic continuous scale which allows to rate 11 chemosensory perceptions from ‘not perceived’ to ‘strongest perception imaginable’ (numeric range: 0-1,000). These measurements will later be sorted according to olfactory perceptions (e.g. smell), annoyance/disgust or sensory irritation (e.g., tingling, nasal irritation, eye irritation etc.).

##### 2.3.2.2 Hedonic

Similar to the LMS, participants rated the perceived air pleasantness on a quasi-logarithmic continuous Labeled Hedonic Scale (LHS) [67] ranging from ‘most unpleasant imaginable’ to ‘most pleasant imaginable’ (numeric range: 0-1,000 with 500 being neutral).

##### 2.3.2.3 SPES

The questionnaire of acute health symptoms (SPES) [68], which was extended by Seeber et al., [69] and van Thriel et al., [48] to include specific chemosensory symptoms, comprises of 29 items (e.g., burning eyes, bad taste in the mouth, cough) that are rated on a scale ranging from 0 (not at all) to 5 (very strong). The items are later averaged into 7 subscales, i.e., olfactory symptoms, taste disturbances, respiratory symptoms, irritative symptoms, nasal irritation, eye irritation, unspecific symptoms. For the current study, taste disturbances and respiratory symptoms were not of interest.

##### 2.3.2.4 Dimensions of Acute Well-Being

Participants rated their acute well-being on a scale from 1 to 6 with opposing descriptions: relaxed – tense, awake – tired, complaints – no complaints, annoyed – not annoyed (see [70]).

##### 2.3.2.5 Working Memory Performance

During the exposure, participants performed two versions of the n-back task, each two times (see Table 2). The n-back is a continuous working memory task in which the participant is presented with a series of 150 images of objects on a screen. Each image is shown for 1500 ms with an inter-stimulus-interval of 1500 ms. During the presentation, participants rest the index finger of their dominant hand on a button that registers whenever the finger is lifted. The participants’ task is to respond by lifting their index finger and pressing a button next to their rest-button whenever the image currently being shown matches the penultimate (2-back) or third-to-last (3-back) image. After each press the participant returns the index finger to the rest-button. Forty-five of the 150 trials were target trials. For the analysis, the sensitivity index *d’* was calculated as follows:

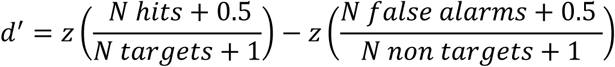

To handle extreme proportions of correct trials 0.5 was added to each cell of the contingency table.

##### 2.3.2.6 Heart Rate

Participants were equipped with a heart rate monitor (FAROS-EKG, Bittium Biosignals) that recorded heart beats throughout the experiment with a sampling frequency of 500 Hz. Analysis time windows were marked online, corresponding to the n-back task period and resting period in which they were asked to sit still and relax (see Table 2). Offline, the HR and HRV were calculated using the open source HRVTool toolbox [71] in Matlab (R2021b, The Math-Works Inc, Natick, USA). This toolbox was additionally used to calculate frequency-domain measurements using the fast fourier transform, i.e., HF (0.15 – 0.4 Hz), and low frequency power (LF; 0.04 – 0.15 Hz) in percentage of total HRV (minus very low frequencies), as well as their ratio LF/HF. As HF and LF are inversely proportional, we will only report HF and LF/HF in the result section. Furthermore, time-domain measurements, i.e., RMSS as well as SDNN were estimated as has been done by Kumpulainen et al. [47]. The focus of the analysis was how the heart rate parameters change from the last minute of the 3-back task (presumed stress induction) and throughout the following 7-minute resting period in 1-minute intervals. Thus, the aim of this analysis was to assess stress recovery over time and how the exposure might interact with the respective heart rate parameters.

### 2.4 Protocol

#### 2.4.1 Training Day

To take part in the main experiment, participants were first invited to undergo a mandatory 2 h training day (either from 9 to 12 am or from 1 to 4 pm). The dates for the main experiment were only scheduled after successful participation in the training. In case the participants were not invited to take part in the main experiment, they were reimbursed for the training with 36 € or university credits.

Participants were shown the ExpoLab and were instructed on how to correctly use the sliders and keyboard to perform the ratings and n-back tasks. Moreover, the nasal lavage, NIOX VERO® measurements, as well as keratography measurements were practiced.

Furthermore, participants were examined using the Sniffin’ Sticks test battery (Burghart Messtechnik GmbH, Wedel, Germany). This standardized smell test comprises three subtests. The identification test consists of 16 different odors that are presented one after the other and must be recognized by the participant from 4 alternative options. The threshold test consists of 16 triplets, each of which is compared with the aim of recognizing the odor n-butanol. The concentration is slowly adjusted in single staircase, 3-alternative forced choice procedure so that the individual odor threshold can be assessed. The discrimination test also contains 16 triplets, which are compared with the aim of recognizing the smell of one sample that differs from the other two. This assesses the test subject’s ability to differentiate between smells.

#### 2.4.2 Experimental Days

Each experimental day followed the same structure (Table 2) and lasted 4.5 hours (either 8 am to 12:30 pm or 11:30 am to 4 pm). Two participants took part at once in parallel. First, participants completed the pre-measurement, i.e., FeNO, keratography, nasal lavage. They were equipped with the ECG belt as well as an EEG cap (data will be published separately). Participants entered the ExpoLab at either 9:20 am or 12:50 pm. Throughout the exposure, participants completed the ratings three times, the n-back tasks twice and resting period four times. Halfway through the exposure, participants could take a break inside the lab and use the restroom or eat a snack (care was taken to select snack options without a strong odor). Each exposure lasted for 2 hours. After the exposure, participants completed the post-measurements.

## 3 Statistical Analysis

The statistical analysis included multi-factorial Analyses of Variance with repeated measures (rmANOVA) conducted in JASP (version 0.95.4), if assumptions were met. In case the assumption of sphericity was violated, the Greenhouse-Geisser (GG) correction was applied. If a parametric method could not be used (e.g., categorical scale or violation of normality), an ANOVA-type analysis was performed using the MANOVA.RM package [72] in R 4.0.3 [73], using RStudio [74]. This analysis is based on a resampling-approach and is considered robust to the breaches of assumptions generally presumed for ANOVAs [75]. Either paired *t*-tests (in JASP) or Wilcoxon signed-rank tests (rstatix package [76] in R), were used to perform post-hoc tests. The level of significance for all statistical tests was set to 0.05. Multiple comparison adjustments were applied using the Bonferroni-Holm method.

All ANOVAs included the factors ‘exposure’ (Table 3). The factor ‘exposure’ consists of three levels corresponding to the exposure conditions: control, GV I and GV II. Further, some analyses contained the factor ‘time’. For the perceptual ratings the factor ‘time’ related to the 3-fold repeated ratings (e.g., hedonic, LMS, etc.). For the changes in heart rate the factor ‘time’ covered the last minute of 3-back task and 7 minutes of the resting period in 1-minute increments (total 8 minutes). To ensure interpretable HRV results within the framework of the current study, only interactions including the factor time are considered for the current manuscript. For transparency reasons, all significant results are reported, yet post-hoc tests are only conducted if the factor time is included in the interaction with any significant main effect. The HRV changes are further split into 1^st^ and 2^nd^ half of the experiment, corresponding to the 2^nd^ and 4^th^ rest periods and respective preceding 3-back tasks (Table 2). Similarly, the 1^st^ and 2^nd^ block of n-back trials were analyzed using the factor ‘halves’. The pre- and post-measurements targeting sensory irritation are expressed as percentage change from the pre- to the post-measurement as follows:

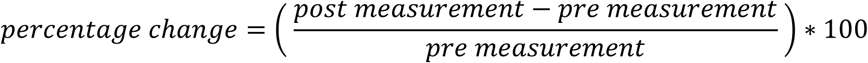

Thus, positive values reflect an increase of physiological measures of sensory and upper airway irritation.

**Table 3.**
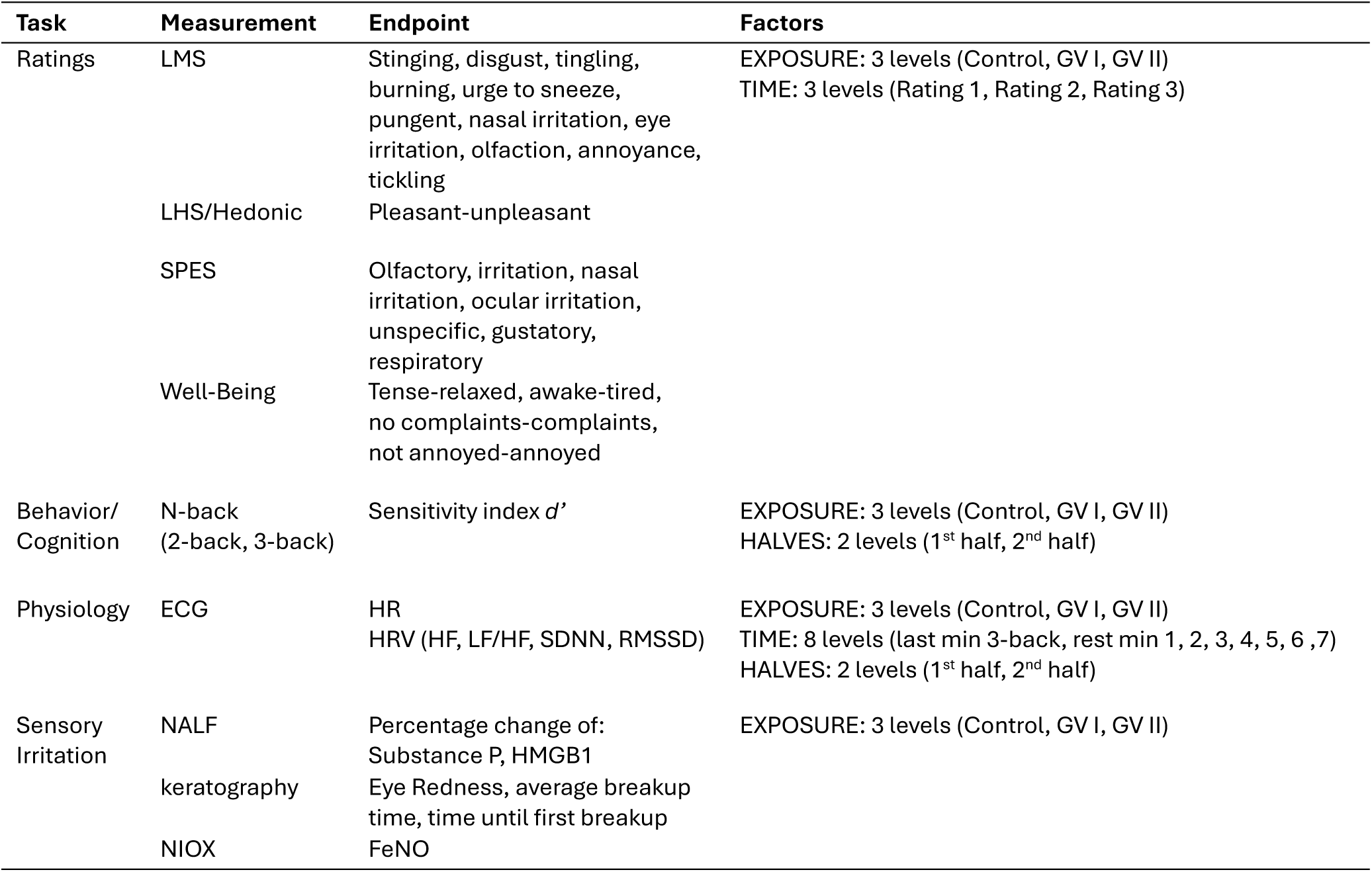
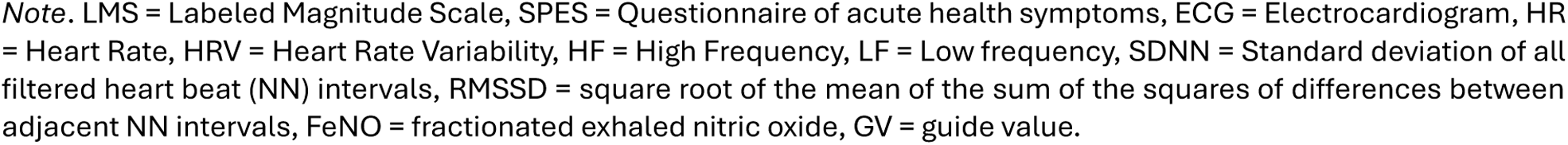
Overview over measurement endpoints of each task and respective factor structure for the Analysis Of Variance(-type) analyses.

| Task | Measurement | Endpoint | Factors |
| --- | --- | --- | --- |
| Ratings | LMS | Stinging, disgust, tingling, burning, urge to sneeze, pungent, nasal irritation, eye irritation, olfaction, annoyance, tickling | EXPOSURE: 3 levels (Control, GV I, GV II)<br>TIME: 3 levels (Rating 1, Rating 2, Rating 3) |
|  | LHS/Hedonic | Pleasant-unpleasant |  |
|  | SPES | Olfactory, irritation, nasal irritation, ocular irritation, unspecific, gustatory, respiratory |  |
|  | Well-Being | Tense-relaxed, awake-tired, no complaints-complaints, not annoyed-annoyed |  |
| Behavior/<br>Cognition | N-back<br>(2-back, 3-back) | Sensitivity index $d'$ | EXPOSURE: 3 levels (Control, GV I, GV II)<br>HALVES: 2 levels (1<sup>st</sup> half, 2<sup>nd</sup> half) |
| Physiology | ECG | HR<br>HRV (HF, LF/HF, SDNN, RMSSD) | EXPOSURE: 3 levels (Control, GV I, GV II)<br>TIME: 8 levels (last min 3-back, rest min 1, 2, 3, 4, 5, 6, 7)<br>HALVES: 2 levels (1<sup>st</sup> half, 2<sup>nd</sup> half) |
| Sensory<br>Irritation | NALF | Percentage change of:<br>Substance P, HMGB1 | EXPOSURE: 3 levels (Control, GV I, GV II) |
|  | keratography | Eye Redness, average breakup time, time until first breakup |  |
|  | NIOX | FeNO |  |

Note. LMS = Labeled Magnitude Scale, SPES = Questionnaire of acute health symptoms, ECG = Electrocardiogram, HR = Heart Rate, HRV = Heart Rate Variability, HF = High Frequency, LF = Low frequency, SDNN = Standard deviation of all filtered heart beat (NN) intervals, RMSSD = square root of the mean of the sum of the squares of differences between adjacent NN intervals, FeNO = fractionated exhaled nitric oxide, GV = guide value.

**Table 4.** Significant post-hoc paired t-tests or Wilcoxon signed-rank tests of heart rate parameter changes over time.

| Comparison | HR |  |  | HF |  |  | LF/HF |  |  | SDNN |  |  |
| --- | --- | --- | --- | --- | --- | --- | --- | --- | --- | --- | --- | --- |
|  | *t* | *p* | *d* | *z* | *p* | *d* | *z* | *p* | *r* | *z* | *p* | *r* |
| Last min 3-back |  |  |  |  |  |  |  |  |  |  |  |  |
| 1<sup>st</sup> min rest | 8.03 | <0.001 | 0.43 | 1.52 | 0.004 | 0.29 | 1.28 | 0.036 | 0.24 | 2.58 | <0.001 | 0.49 |
| 2<sup>nd</sup> min rest | 7.68 | <0.001 | 0.46 | 2.13 | <0.001 | 0.40 | 2.15 | <0.001 | 0.41 | 3.01 | <0.001 | 0.57 |
| 3<sup>rd</sup> min rest | 6.21 | <0.001 | 0.43 | 2.28 | <0.001 | 0.43 | 2.13 | <0.001 | 0.40 | 3.12 | <0.001 | 0.59 |
| 4<sup>th</sup> min rest | 6.32 | <0.001 | 0.40 | 1.91 | <0.001 | 0.36 | 1.62 | 0.002 | 0.31 | 3.24 | <0.001 | 0.61 |
| 5<sup>th</sup> min rest | 5.89 | <0.001 | 0.41 | 2.28 | <0.001 | 0.43 | 2.07 | <0.001 | 0.39 | 3.17 | <0.001 | 0.60 |
| 6<sup>th</sup> min rest | 6.48 | <0.001 | 0.46 | 2.02 | <0.001 | 0.38 | 1.81 | <0.001 | 0.34 | 3.25 | <0.001 | 0.61 |
| 7<sup>th</sup> min rest | 4.81 | 0.001 | 0.35 | 1.59 | 0.002 | 0.30 | 1.57 | 0.003 | 0.30 | 3.72 | <0.001 | 0.51 |

Note. HR = Heart rate, HF = High Frequency, LF = Low frequency, SDNN = Standard deviation of all filtered heart beat (NN) intervals, RMSSD = square root of the mean of the sum of the squares of differences between adjacent NN intervals.

## 4 Results

For a more concise presentation of the relevant results, only significant results will be presented in the main manuscript. For a complete list of results, please refer to the online OSF repository containing all (partly original partly preprocessed due to sensitive information) anonymized data, R and matlab scripts and JASP files (will be made available for reviewers and publicly available after acceptance [here](https://osf.io/kstue/overview?view_only=246c2e54b00d47edaa147bb312451484)). Effects that show changes over time will be displayed in blue while changes evoked by the different exposure scenarios will be displayed in yellow-orange tones.

### 4.1 Rating Results

#### 4.1.1 LMS Intensity Ratings

Ratings of eye irritation intensities were significantly influenced by measurement timepoint which corresponds to time in the ExpoLab, *ATS* = 7.44, *p* < 0.001, *η_p_^2^* = 0.19 (Figure 1 A). Exposure concentration as well as the interaction were not significant. Post-hoc Wilcoxon signed-rank tests revealed an increase in eye irritation intensity with significant differences between the first and second (*z* = 2.21, *p* < 0.001, *r* = 0.39) as well as first and third time point, *z* = 2.50, *p* < 0.001, *r* = 0.44.

**Figure 1.**
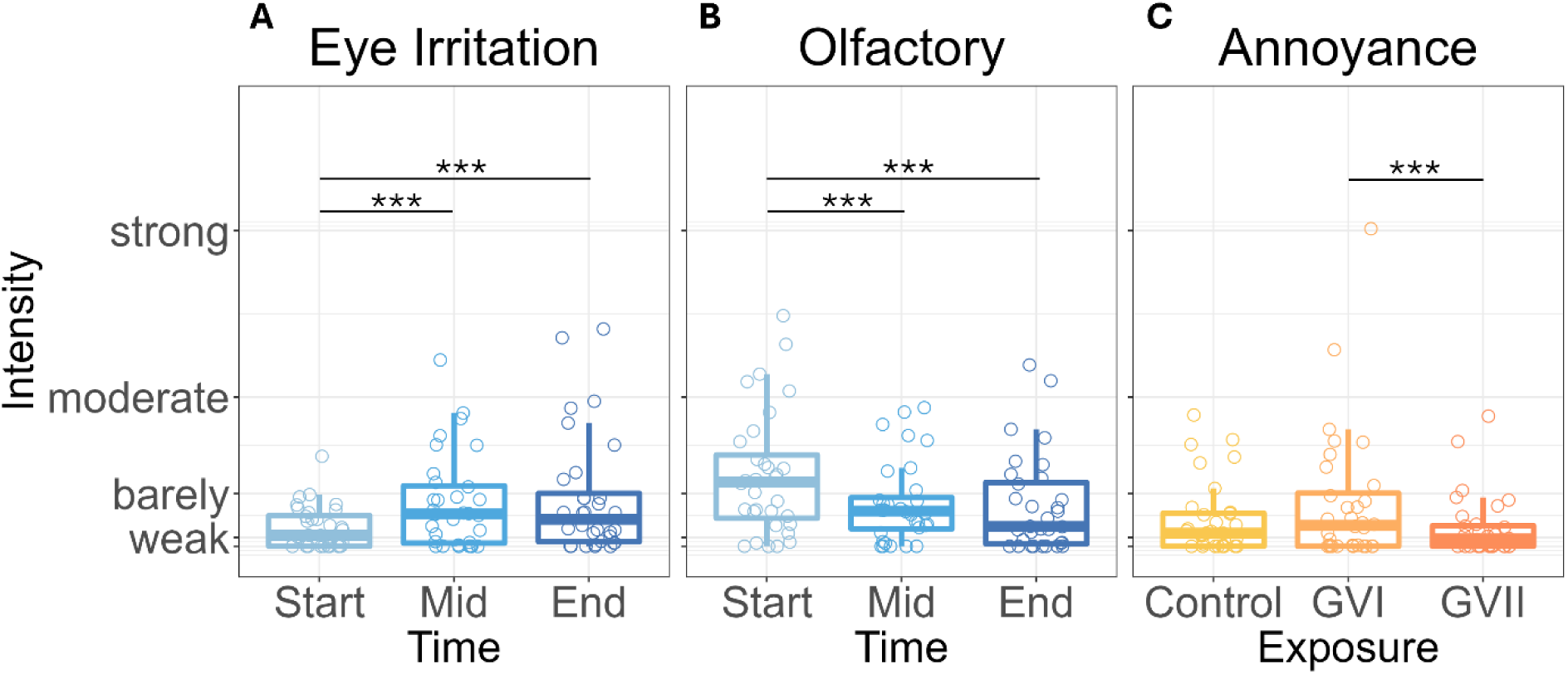
Significant intensity ratings based on the Labeled Magnitude Scale (LMS) of eye irritation (A), olfactory (B) and annoyance (C). *** p < 0.001.

Similar effects were shown for olfactory intensity ratings which were also significantly influenced by time, *ATS* = 8.18, *p* < 0.001, *η_p_^2^* = 0.21. Again, neither the effect of exposure scenarios nor the interaction was significant. Post-hoc Wilcoxon signed-rank tests revealed a significant decrease in olfactory intensity ratings between the first and second (*z* = 2.15, *p* < 0.001, *r* = 0.38) as well as first and third time point, *z* = 2.12, *p* < 0.001, *r* = 0.38 (Figure 1 B).

Annoyance, however, was influenced by exposure scenario as revealed by a significant main effect for ‘exposure’, *ATS* = 4.16, *p* = 0.02, *η_p_^2^* = 0.12. Post-hoc Wilcoxon signed-rank tests showed that participants rated the annoyance intensity as significantly stronger in the GV I compared to the GV II exposure condition, *z* = 2.09, *p* < 0.001, *r* = 0.37 (Figure 1 C).

Trigeminal perceptions of tickling, stinging, tingling, burning, urge to sneeze, pungent, nasal irritation (trigeminal irritations) or ratings of disgust were neither influenced by measurement time point nor exposure condition.

#### 4.1.2 Ratings of Hedonics

LHS ratings were significantly influenced by exposure conditions, evident by a significant main effect, *ATS* = 3.68, *p* = 0.03, *η_p_^2^* = 0.23. Post-hoc Wilcoxon signed-rank tests showed significantly lower hedonic ratings for the GV I compared to the control (*z* = 1.32, *p* = 0.04, *r* = 0.23) as well as GV II exposure scenarios, *z* = 1.42, *p* = 0.04, *r* = 0.25 (Figure 2).

**Figure 2.**
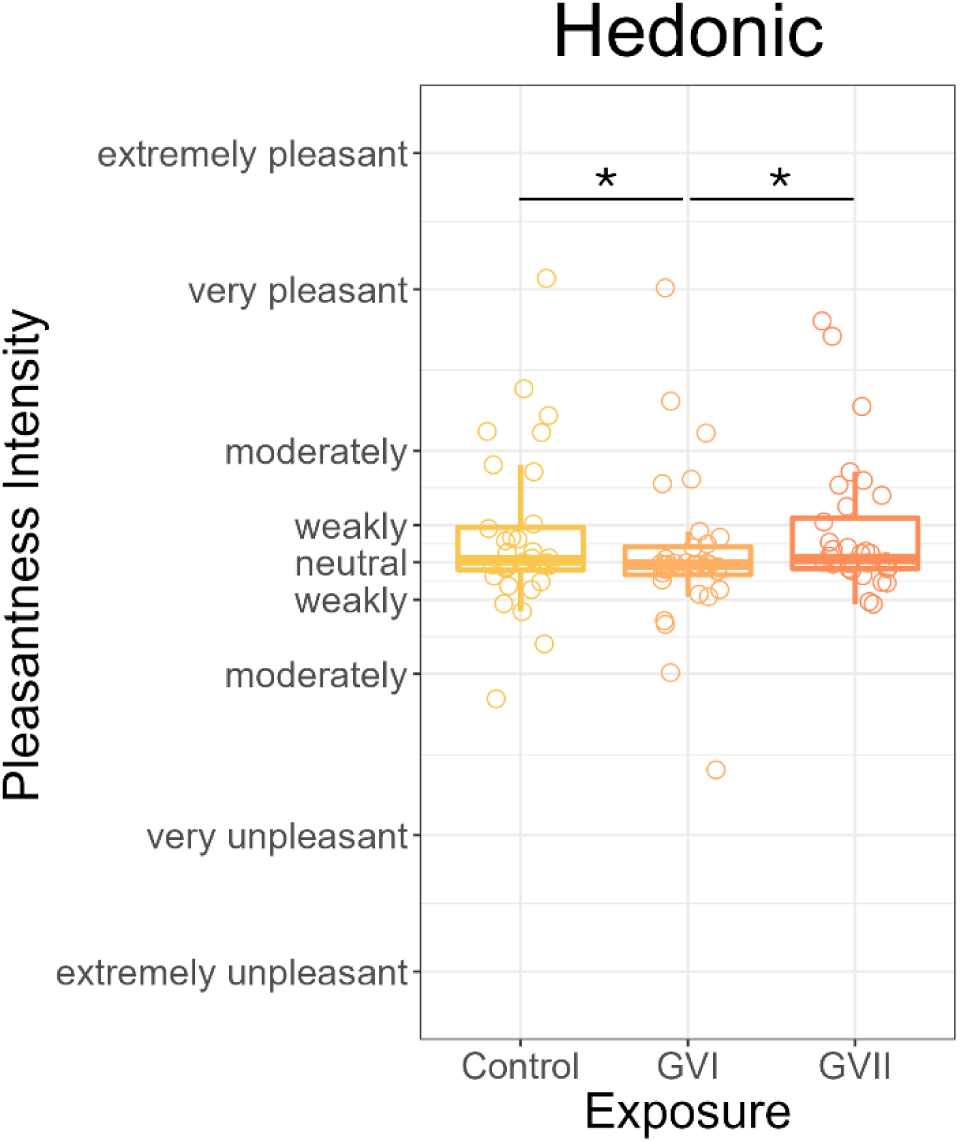
Hedonic ratings with significant differences between the guide value (GV) I and control as well as GV II condition. * p < 0.05.

#### 4.1.3 SPES Symptom Questionnaire

The ANOVA-type statistic revealed that nasal irritations was influenced by time in the ExpoLab, *ATS* = 3.97, *p* = 0.02, *η_p_^2^* = 0.11. This factor, however, also significantly interacted with the factor ‘exposure’ scenario, *ATS* = 3.22, *p* = 0.02, *η_p_^2^* = 0.09. Post-hoc Wilcoxon signed-rank tests showed an increase over the experimental time with a significant difference between the last measurement point of the GV II exposure condition and the first measurement point of the control (*z* = 3.16, *p* = 0.002, *r* = 0.56), GV I (*z* = 3.04, *p* = 0.004, *r* = 0.54), and GVII condition (*z* = 2.27, *p* = 0.05, *r* = 0.40) as well as the last measurement point of the control condition, *z* = 2.35, *p* = 0.03, *r* = 0.42 (Figure 3 A). Moreover, the nasal irritation ratings of the second measurement point of the control condition were also significantly higher than the first measurement point of the control condition (*z* = 1.98, *p* = 0.03, *r* = 0.35).

**Figure 3.**
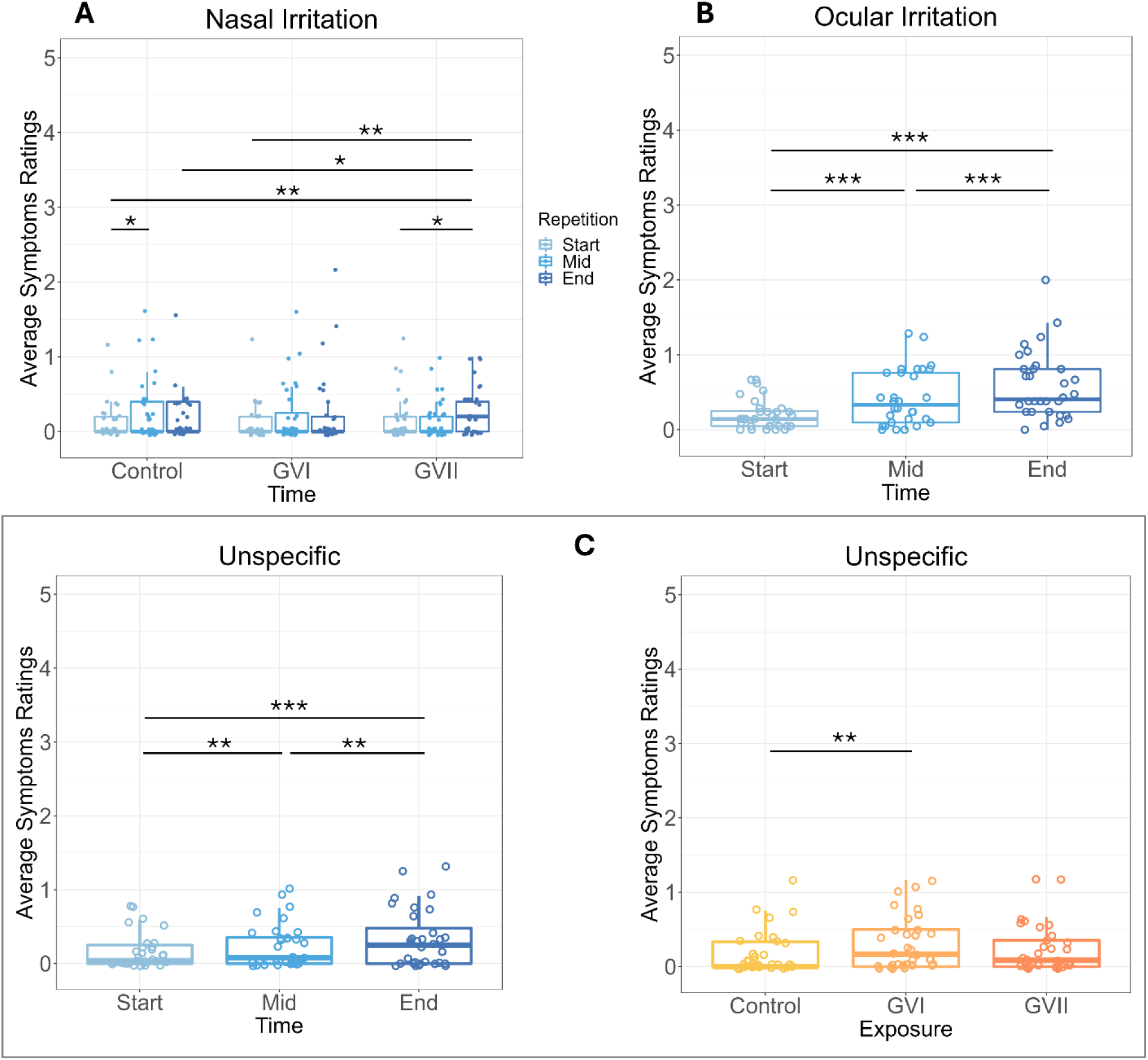
Ratings of significant SPES symptoms that are averaged from sub items regarding specific health symptoms, i.e., nasal irritation (A), ocular irritation (B), and unspecific symptoms (C). * p < 0.05, ** p < 0.01, *** p < 0.001.

Ocular irritation also changed over time as shown by a significant main effect, *ATS* = 26.25, *p* = 0.001, *η_p_^2^* = 0.56 (Figure 3 B). Yet, neither the factor ‘exposure’ nor the interaction was significant. A continuous increase in ocular irritation was supported by post-hoc Wilcoxon signed-rank tests which showed significant differences between all three measurement time points (start vs middle: *z* = 2.98, *p* < 0.001, *r* = 0.527; start vs end: *z* = 4.04, *p* < 0.001, *r* = 0.72; middle vs end: *z* = 2.70, *p* < 0.001, *r* = 0.48).

Unspecific symptoms were influenced by exposure time as well as exposure concentration as evident by a main effect of exposure scenario (*ATS* = 3.58, *p* = 0.03, *η_p_^2^* = 0.10) as well as measurement time point, *ATS* = 13.72, *p* < 0.001, *η_p_^2^* = 0.31 (Figure 3 C). However, no interaction was shown between these factors. Post-hoc Wilcoxon signed-rank tests showed that the symptoms were rated as significantly stronger in the GV I compared to the control condition, *z* = 2.08, *p* = 0.006, *r* = 0.37. Moreover, the ratings increased over the course of the experiment with significant differences between the first and second, *z* = 1.75, *p* = 0.003, *r* = 0.31, first and third (*z* = 2.67, *p* < 0.001, *r* = 0.47) as well as second and last measurement time point, *z* = 1.81, *p* = 0.003, *r* = 0.32.

Olfactory, gustatory, respiratory, or generally irritative symptoms were not influenced by either factor.

#### 4.1.4 Well-Being Results

Participants became less relaxed and more tense throughout the experiment shown by a significant main effect for time, *ATS* = 10.30, *p* < 0.001, *η_p_^2^* = 0.25. Consecutive post-hoc Wilcoxon signed-rank tests showed a significant increase from the start compared to the middle (*z* = 1.09, *p* = 0.039, *r* = 0.19), as well as end of the experiment, *z* = 2.39, *p* < 0.001, *r* = 0.42 and between the middle and final measurement point, *z* = 1.71, *p* = 0.003, *r* = 0.30 (Figure 4 A).

**Figure 4.**
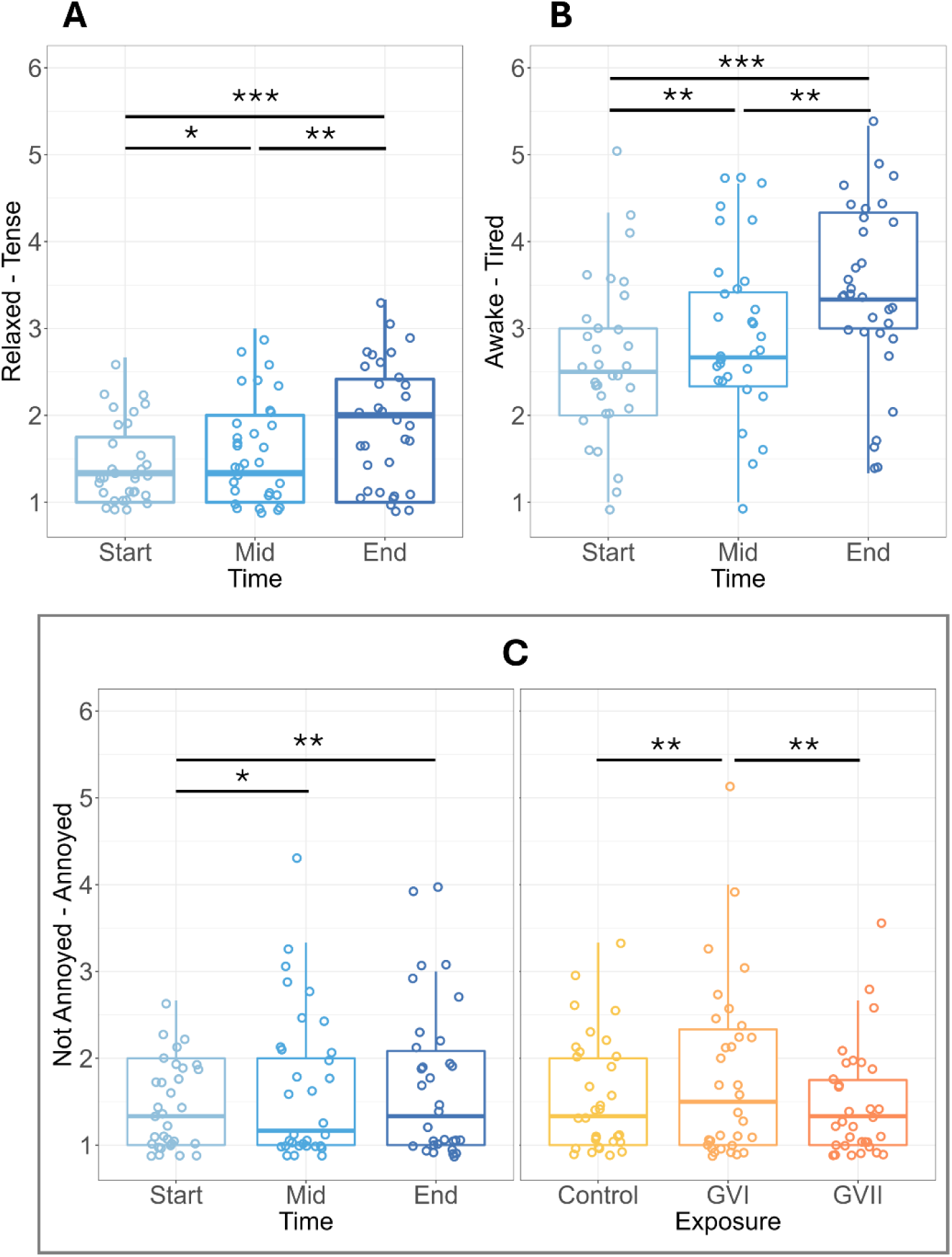
Significant well-being results that indicate the effect of time on relaxed-tense (A), awake-tired (B) and additionally an effect of exposure on not annoyed-annoyed (C). * p < 0.05, ** p < 0.01, *** p < 0.001.

Similarly, participants became more tired over the course of the experiment, indicated by a significant main effect for time, *ATS* = 11.5, *p* < 0.001, *η_p_^2^* = 0.27 (Figure 4 B). Again, all time points differed significantly from each other, start vs. middle: *z* = 1.65, *p* = 0.006, *r* = 0.29; start vs. end: *z* = 2.74, *p* < 0.001, *r* = 0.49, middle vs. end: *z* = 2.15, *p* = 0.001, *r* = 0.38.

Mirroring the effect of the LMS, annoyance showed a significant main effect for exposure scenario (*ATS* = 4.29, *p* = 0.03, *η_p_^2^* = 0.12) with significant post-hoc comparisons showing higher annoyance ratings for the GV I compared to the control (*z* = 1.60, *p* = 0.006, *r* = 0.28) as well as GV II condition, *z* = 1.77, *p* = 0.003, *r* = 0.31. Additionally, the ANOVA-type statistic also indicated a significant effect of time, *ATS* = 3.86, *p* = 0.04, *η_p_^2^* = 0.11. Post-hoc Wilcoxon signed-rank tests revealed an increase in annoyance ratings with significant differences between the first and second (*z* = 1.07, *p* = 0.03, *r* = 0.19) as well as third measurement time point, *z* = 1.52, *p* = 0.006, *r* = 0.27 (Figure 4 C).

General health complaints initially also showed a significant main effect for the factor time, yet none of the post-hoc comparisons remained significant after correction for multiple comparisons.

### 4.2 N-back Results

For the analysis, 2 participants were excluded due to incorrect task performance, i.e., incorrect button use which results in unevaluable performance values. Neither in the 2-back (all p > 0.22) nor the 3-back (all p > 0.47) tasks were the *d*’ affected by the exposure scenario or halves. For more details of the statistical results, please refer to Supplement 4 and the OSF repository.

### 4.3 ECG Results

For the heart rate, the rmANOVA showed significant main effects for factor ‘time’, *F_GG_*(3.85,104.0) = 17.08, *p* < 0.001, *η_p_^2^* = 0.39, ‘exposure’, *F_GG_* (1.97,53.15) = 3.44, *p* = 0.04, *η_p_^2^* = 0.11, and ‘halves’, *F_GG_* (1,27) = 18.31, *p* < 0.001, *η_p_^2^* = 0.40 (Figure 5 A). Post-hoc paired *t*-tests showed significantly lower HR values for GV I compared to GV II, *t*(27) = −2.56, *p* = 0.05, *Cohen’s d* = −0.38. Furthermore, participants had significantly higher HR in the second compared to the first half of the experiment, *t*(27) = 4.21, *p* < 0.001, *Cohen’s d* = 0.47. Moreover, investigating the change over time revealed significantly higher HR for the last minute of the 3-back task compared to all other time points (*Table 5*).

**Figure 5.**
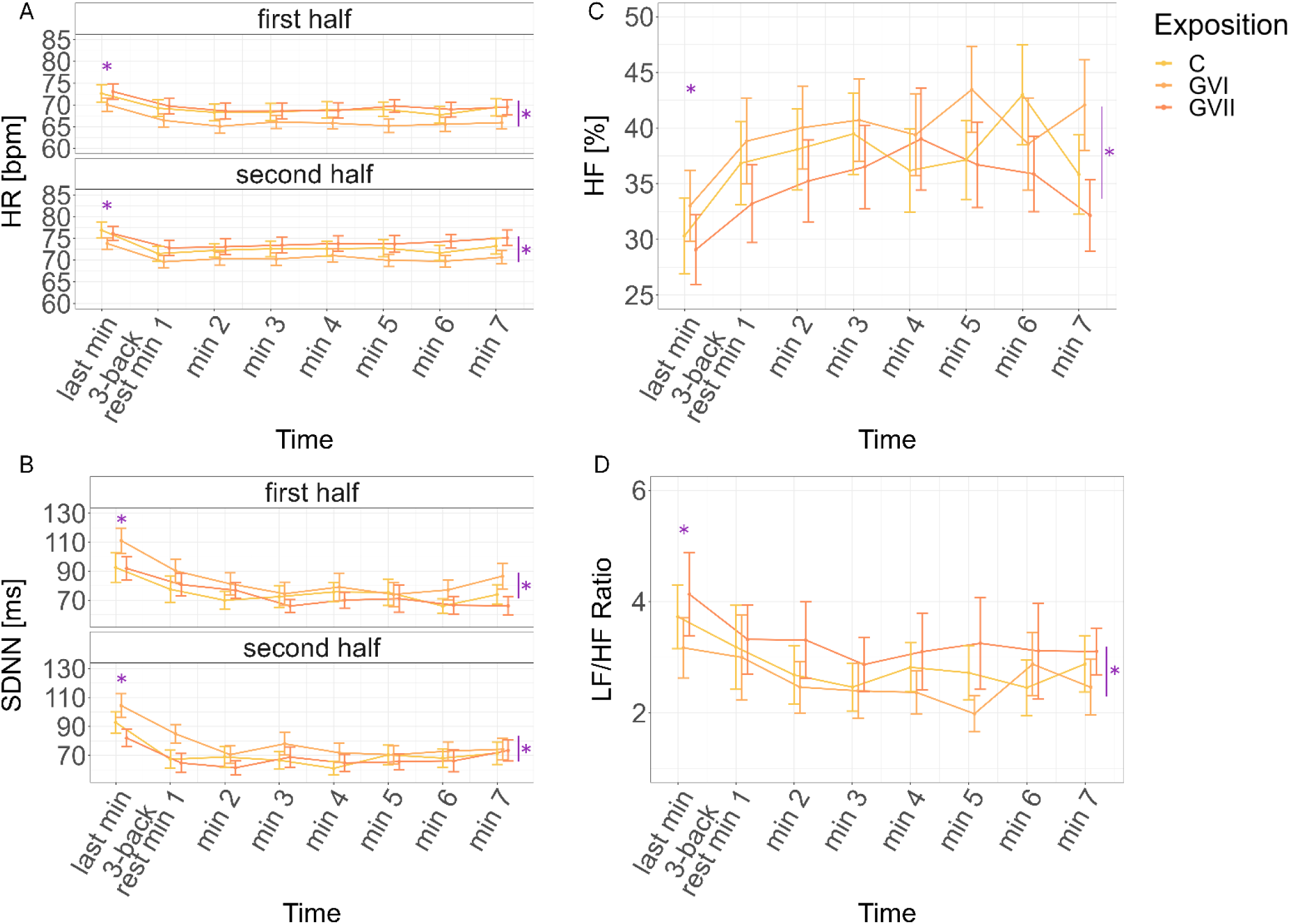
Changes in (A) heart rate (HR) and features of heart rate variations, i.e., (B) standard deviation of normal-to-normal heartbeat intervals (SDNN), (C) relative high frequency power (HF) of the HRV as well as (D) low frequency and HF ratio (LF/HR). Purple asterisk above the first time point indicates a significant difference of this to every other time point. Asterisks on the right-hand side of each plot indicate a significant difference between exposure conditions. A split of plots into first and second half connotes a significant main effect for experimental halves. Error bars depict standard errors.

SDNN was affected by time (*ATS* = 18.70, *p* < 0.001, *η_p_^2^* = 0.41), exposure (*ATS* = 3.09, *p* = 0.05, *η_p_^2^* = 0.10), and differed significantly across the two experimental halves, *ATS* = 4.8, *p* = 0.03, *η_p_^2^* = 0.15 (Figure 5 B). Post-hoc Wilcoxon signed-rank tests indicated that SDNN were significantly higher for the GV I compared to control (*z* = 1.04, *p* < 0.001, *r* = 0.20) as well as to GV II values, *z* = 1.19, *p* < 0.001, *r* = 0.22. Further, SDNN values for the 1^st^ half were significantly higher than for the 2^nd^ half, *z* = 0.80, *p* < 0.001, *r* = 0.15. The SDNN values changed over time in that all time points of the resting phase were significantly lower than the last minute of the 3-back task (*Table 5*).

RMSSD was influenced by exposure (*ATS* = 3.12, *p* = 0.05, *η_p_^2^* = 0.10) in that values for GV I were significantly higher than for the control (*z* = 1.20, *p* < 0.001, *r* = 0.23) as well as GV II condition, *z* = 1.43, *p* < 0.001, *r* = 0.27. No other factor, importantly ‘time’, had an influence on RMSSD.

HF was influenced by exposure (*ATS* = 3.66, *p* = 0.03, *η_p_^2^* = 0.12) and time, *ATS* = 5.91, *p* < 0.001, *η_p_^2^* = 0.18 (Figure 5 C). Post-hoc Wilcoxon signed-rank tests showed the highest HF values for GV I compared to the control (*z* = 0.56, *p* = 0.02, *r* = 0.11) as well as the GV II condition, *z* = 1.16, *p* < 0.001, *r* = 0.22. Furthermore, GV II showed the lowest values with significantly lower HF power compared to the control condition, *z* = 0.63, *p* = 0.02, *r* = 0.12. Again, the time effect resolved in significantly higher HF values in all time points of the rest period compared to the last minute of the 3-back task (*Table 5*).

Lastly, the LF/HF ratio was influenced by time (*ATS* = 3.95, *p* < 0.001, *η_p_^2^* = 0.13) and exposure, *ATS* = 3.69, *p* = 0.03, *η_p_^2^* = 0.12 (Figure 5 D). Post-hoc Wilcoxon signed-rank tests showed significantly lower values for the GV I compared to the control (*z* = 0.56, *p* = 0.03, *r* = 0.11) as well as GV II condition, *z* = 1.12, *p* < 0.001, *r* = 0.21. A further difference was indicated between the GV II and control condition, indicating the highest values for the GV II condition, *z* = 0.69, *p* = 0.011, *r* = 0.13. Again, the time effect manifests in a difference between the last minute of the 3-back task and all other time points (*Table 5*).

### 4.4 Physiological Irritation Results

The NALF results are based on 28 data sets as 5 participants were excluded (due to e.g., failure to participate in all sessions, swallowing of the liquid, detection difficulties during the assay, etc.). Neither the percentage change of the concentrations for Substance P (*p* = 0.14) nor HMGB1 (*p* = 0.42) were influenced by exposure. However, it needs to be mentioned that for HMGB1 41 data points were substituted by the limit of quantitation (BG = 0.89 ng/mL) as they fell below this limit. Keratography measurements did not indicate an exposure-related increase of objective eye irritation, i.e., eye redness (*p* = 0.39) or tear film stability (BUT1: *p* = 0.23, BUM: *p* = 0.16). As for the FeNO results, the exposure did not influence the measurements (*p* = 0.54).

## 5 Discussion

The goal of this study was to simulate a demanding workday-like situation to explore the effects of exposure to pinewood VOCs in concentrations between the GV I and GV II (as measured by α-pinene and 3-caren) compared to clean air. We investigated if the emissions (1) are perceived, (2) rated as pleasant or unpleasant/annoying, (3) induce relaxation vs. stress on the perceptual or physiological level and/or (4) induce measurable indicators of sensory irritation. Ratings indicate that the time of the exposure sessions, independent of the condition, had a tiresome effect on the participants (effects visually depicted in blue, e.g., SPES ratings of awake, LMS ratings of eye irritation). Thus, we can assume that the goal of creating a 2 h workday type situation was successful. Still, it is important to note that most effects of exposure, even if statistically significant, are to be considered as small changes and physiologically unobtrusive.

### 5.1 Olfactory Perception of Pinewood VOCs

Based on the literature, it was expected that if participants perceived the pinewood odor in the whole-body exposure, they would rate it as pleasant [21], [41], [77] due to the generally positive attitude towards wood [78], [79]. Reported olfactory detection thresholds for α-pinene differ vastly from 1.3 mg/m^3^ [80], to 23 mg/m^3^ [81] to even 107 mg/m^3^ [82], depending on the assessment method and enantiomer. It is known that different assessment methods result in different thresholds. A whole-body exposure might lead to different experiences and derived thresholds than olfactometry results [83], [84], [85]. Thus, it was difficult to predict whether participants would be able to detect wood VOCs in concentrations of the GV values. Furthermore, there are more odor-active compounds in pinewood samples (e.g., octenal, *(E)-*non-2-enal, or hexanal) that might have fallen above the olfactory detection threshold in the current study [86], [87]. Therefore, it was possible that participants could have experienced an olfactory perception of the wood emission in our experiments.

Ratings of olfactory intensity (LMS) show that participants initially reported the perception of an odor which then faded due to habituation and/or adaptation of the olfactory system [21], [88]. Nevertheless, we did not find a main effect for exposure (meaning olfactory perceptions were also present in the control condition), nor an interaction of exposure and time. Thus, we cannot conclude that the participants could successfully differentiate between the three exposure conditions in terms of olfactory intensity. Thus, unusual and purely ExpoLab-related odors or the air conditioning could be responsible.

In terms of pleasantness, our results indicate that the pinewood odor was not evaluated and rated to be more or less pleasant than the control condition. Statistically, the GV I condition was rated to be slightly less pleasant than the other conditions. Surprisingly, these differences occurred without any differences in odor intensity. This pattern aligns with the annoyance ratings (LMS and SPES) which indicate the highest annoyance ratings for the GV I condition compared to the other two. However, numerically, all conditions centered around ‘neutral’. This might reflect that the participants simply did not perceive the odors as indicated by the above-mentioned olfactory intensity ratings. Non-sensory factors might also contribute to these subtitle differences. However, previous studies using higher concentrations of pinewood showed that the odor hedonics were rated as rather neutral or slightly pleasant with large inter-individual difference [37], [86], [87], [89]. Contrarily, [77] showed that among different wood types, pinewood was rated to be the most pleasant. These studies differ in how the odor was presented and these methods, again, differ from the continuous whole-body exposure of the current study. Interestingly, Gminski et al., [21] could not show concentration-dependent differences in intensity nor pleasantness in a whole-body exposure for pinewood emissions either. These mixed results suggest methodological dependencies as well as generally high inter-individual differences in the hedonic evaluation of pinewood.

Inter-individual differences may be the most likely underlying reason for the concentration-independent hedonic profile in the current study. Bontempi et al., [90] showed an interaction between hedonic ratings and concentration in a way that pleasant odors tend to become more pleasant whereas unpleasant odors are rated as more unpleasant with increasing concentration. This relationship in turn depends on the individual detection thresholds. Perhaps, part of our sample rated pinewood as more pleasant whereas the other as less pleasant, and the different concentrations (GV I and GV II) affect the hedonic ratings, respectively. Yet it is noteworthy that in the GV II condition the ratings do not fall within the negative range as much as in the other two conditions which would not fit this narrative.

Moreover, studies that suggest positive effects of woods oftentimes use visual features [47]. In a previous study, we could demonstrate that even if participants were prompted to only rate olfactory pleasantness of pinewood odors, concurrently shown pictures of either wood or non-wood patterns influenced the ratings significantly. This suggests a holistic processing of multisensory percepts [89]. In a recent study Jyske and colleagues [41] demonstrated that in a whole-body exposure participants did perceive wood scents as more pleasant than no scent, but virtual wall displays of wood textures massively influenced the positive effect on stress and restoration. Song et al., [39] also demonstrated a positive effect of visual, olfactory and especially combined nature stimuli on HRV but also subjective ratings of relaxation and comfort. Thus, visual perceptions are an important modulating factor when judging the psychophysiological effects of wood VOCs.

All in all, the large inter-individual differences, the diverse reports of pleasantness regarding wood odors (or α-pinene) as well as the influence of the other senses highlight the challenges for attempts to regulate olfactory experiences of low-level VOC concentrations as usually found in indoor air environments.

### 5.2 Induction of Relaxation vs. Stress

Pinewood VOCs (or individual components such as α-pinene) can induce relaxation. This might be due to terpenes influencing the GABAergic system [91], [92] and/or by increasing positive affect due to the exposure to nature as proposed by the SRT [46] or the ART [51]^i^. It was therefore proposed that if there is a measurable effect of the exposure to pinewood VOCs, participants would (1) feel more relaxed and (2) show a more relaxed bodily state in the rest period after the n-back tasks when exposed to pinewood VOCs compared to the control condition. This effect should be concentration dependent [35].

Subjectively, the ratings of tension-relaxation did not support the hypothesis. While tension increased (relaxation decreased), this effect was not influenced by exposure to VOCs in any way. Thus, the demanding tasks or other non-sensory factors are more likely to cause this overall within session effect of increasing tension.

As objective markers, HR, HF and LF/HF indicated that the induction of stress via the 3-back task was successful and supported the assumption of a consecutive bodily relaxation in the resting period. Whereas RMSSD did not show any response SDNN did show a reversed pattern. High SDNN usually indicates physiological resilience against stress, and we expected to show an increase during the relaxation period. However, SDNN is usually thought to be most effective to be used in a 24 h measurement and it can easily be confounded by changes in respiration patterns [42],[43]. While this reversed pattern might indicate that participants adapted well to the increased task demands, it is difficult to interpret (also see [93] for reversed results indicating an effect of anticipation). We will therefore focus on HR, HF and LF/HF.

Generally, VOC exposure did not interact with the relaxation response of HR, HF and LF/HF per se as no statistically significant interaction between time and exposure was found in any of the recorded parameters. Solely main effects of exposure indicated vertical linear shifts. However, the directions of these effects are in contrast with the hypothesis. HR was higher in the GV II compared to the GV I condition. Higher HR is usually interpreted as stressed state of the body [44]. This indicates a generally less relaxed or higher stress state during the GV II compared to the GV I exposure. HF is used as a surrogate for parasympathetic nervous system activity, as it reflects the activity of the vagus nerve [44]. Thus, higher HF proportions indicate stronger relaxation which, in the current study, was most strongly associated with the GV I condition, followed by the control condition and the lowest proportion was found in the GV II condition. The LF/HF mirrored this pattern. Thus, these parameters suggest the strongest bodily relaxation during the GV I exposure, followed by the control condition and the lowest relaxation during the GV II exposure. Again, the differences are purely based on statistical significance, and the magnitude of these effects is physiologically unobtrusive.

While most studies report evidence in favor of the physiologically relaxing effect of terpene-dominated wood VOCs, others fail to show consistent effects across all parameters. For instance, on the one hand Matsubara and colleagues [94] did show a slowing effect of *Abies sibirica* (Pinaceae) essential oil which also contains a high proportion of terpenes, on HR during a rest period following a task. On the other hand, no effects on the HF proportion could be shown. Further, Ikei and colleagues [34] reported that stimulation with α-pinene also reduced HR and did increase HF. Yet, no effect on the LF/HF ratio was found. While these results were interpreted as a sign of physiological relaxation, participants did not indicate to be more relaxed, only more comfortable. This supports the general assumption that α-pinene might induce relaxation yet contradicts the current results in the form of how it affects the body as well as the perceptions. Moreover, Matsubara and Kawai [95] reported lower LF/HF ratio due to olfactory stimulation with hidden Japanese Cedar panels during a mental arithmetic task but HF proportions were not significantly different from the control exposure. Thus, even studies in support of the original hypothesis fail to yield consistent results, and others even show no effect on HR/V parameters whatsoever [37], [96].

Interestingly, Ojala and colleagues [97] also found signs of a more active sympathetic nervous system as indicated by HRV parameters when participants completed mental work and rested in a wooden vs. a non-wooden room. This was despite a perceived reduction in anxiety. This might be in line with results by Shima, Maeda, et al., [38] who differentiated between different types of arousal evoked by stimulation with different odors: tense vs. energetic. The measured alleged stress response might have been an energetic arousal with similar physiological footprint but opposing psychological effects. This might relate to results by Sorokowska and colleagues [98] showing that odors are processed differently when they are arousing and that this does not necessarily relate to hedonics. Interestingly, they have used pine needle scent as the ‘pleasant arousing’ stimulus which is not in line with the general idea of calming effects of pinewood. Thus, in the current study, the pinewood VOCs might fall into the category of pleasant and energetic arousing yet without a necessary increase in negative tension nor relaxation.

Critically, the missing interaction does not indicate a relaxation or stress-relief booster after the working memory task but rather an overall effect throughout the experiment. Shima, Sakata, et al., [99] stimulated participants with Hinoki cypress scents during work or during rest and could show that only the stimulation during rest increased relaxation while an exposure during work increased negative mood. Thus, the effect was not an induction of relaxation but rather a strengthening of an existing relaxation. Perhaps, participants in the current experiment were overall not relaxed enough due to the experimental setting itself to benefit from the relaxing properties of pinewood VOCs. Future studies should dive deeper into this complex matter, disentangling the physiological markers of arousal as well as effects of stimulation with pinewood VOCs at different time points during work and rest. These results have clear implications for the challenges of regulating odors. The same odor might have opposing effects during work and rest, both occurring in the same room with the same VOC profile in e.g., office buildings.

Circling back to a previous argument, Zhang et al., [40] showed that wooden elements in the experimental room influence HR and HRV in a way that was originally suspected. As no VOCs were measured, we can only conclude that visual features evoked or at least contributed to the recorded effects. This is supported by results in which parts of the wooden elements were painted, thus removing some of the visual features without altering the VOC profile. This is in line with Bamba and Azuma [100] showing psychological stress relief specifically in a room with visual and olfactory wooden features. Olfactory features alone were not different to the control condition. Interestingly, HR, HF, LF/HF were not affected by the olfactory stimulation. Thus, the lack of (visual) context during the VOC exposure in the current study might have influenced how participants evaluate the exposure and respectively how the autonomous nervous system reacts to the stimulation that cannot be certainly categorized as stemming from a natural source without the visual cues. To complicate the matter even more, Sakuragawa and colleagues [101] demonstrated that visual features had a positive effect on the physiology of the observers but only on those, that indicated to like them. This further highlights the effect of inter-individual differences.

Other inter-individual differences e.g., how humans react to psychological stress [102], might contribute to the non-concentration dependent changes in HR and HRV as a result of the n-back task and resting period. This is at least in parts supported by the rather large variance in the HRV data e.g., HF. Research has shown that stress affects the olfactory detection threshold of malodors [61]. Perhaps, a subsample of the participants felt particularly stressed by the n-back task and perceived the odor at lower concentrations. Thus, the individual stress level might have influenced the results in a way that was not expected. Other factors, such as the connectedness to nature might have influenced the individual responses. Demattè and colleagues [33] could show that the individual biophilia degree appears to influence the tactile, auditory, and olfactory evaluation of wooden vs. non-wooden settings.

All in all, the pattern of the data (i.e., non-concentration dependence) as well as the large variation suggest that individual factors that were not recorded as covariates bled into the results. A larger-scale participant sample is needed in future studies to investigate the effect of inter-individual differences, preferences, or stress-responses and relaxation patterns, when exposed to VOCs of terpene-heavy wood odors.

### 5.3 Sensory Irritation

As the exposure concentrations were defined by the GV values, no signs of sensory irritation were expected. This was supported by the pre- vs. post-measurements i.e., NALF, keratography measurements, FeNO. While subjective ratings indicated a slight form of eye irritation (LMS), objective markers such as eye redness or tear film stability were not affected by the experiment nor the experimental conditions. Nasal irritation symptoms as indicated by the SPES questionnaire are the only measure that in fact shows an interaction between time and exposure indicating slightly higher symptoms over time in the GV II condition. However, it needs to be stressed that while this effect was statistically significant, the numeric ratings were still very low, and comparable LMS ratings did not confirm this observation. Care was taken to only invite subjects without acute allergies, however, other factors might have made a sub-group more vulnerable. Suzuki and colleagues [103] demonstrated that especially people with allergies or a high sensitivity to chemicals experience stronger symptoms related to indoor VOCs emitted from building structures. Moreover, Pacharra, Kleinbeck, Schäper, Juran, et al., [60] summarized different exposure experiments and demonstrated that a complex interplay of individual factors (e.g., sex and trait anxiety) markedly influenced ratings of irritation. Thus, in the current study, it is possible that acute mental states, trait-like or demographic factors or even a combination thereof might have influenced the results. Future studies are needed to pinpoint important influential factors to be included in risk assessments of VOCs.

Thus, apart from a slight increase in nasal irritation symptoms, the current exposure study does not suggest any risk for sensory irritation evoked by the VOCs. This is in line with previous studies that did not show any reactions either [81], [104]. Reported thresholds for sensory irritation are much higher [20], and – even at a large cohort scale – no risks could be shown for e.g., asthma development [22]. Thus, current GV seem to protect from irritating effects of pinewood VOC emissions monitored via the concentration of bicyclic terpenes. Especially as the range between the GV I and GV II calls for counteractive measures which will decrease the indoor concentration, diminishing these slight effects. Effects on nasal irritation symptoms should be investigated in future studies to rule out any subtle effects that could not be directly linked to any individual confounding factor in the current study.

### 5.4 Consequences for performance

As there were no strong effects on the perceptual or irritational level it would have been unexpected if working memory performance compromises due to exposure scenarios were found. Unsurprisingly, the performance in the n-back tasks was not affected by the exposure. This is in line with studies showing that even exposures that cause a strong response (e.g., annoyance) only show effects on behavioral performances on a subset of tasks if any [70], [105]. While there are studies that show beneficial effects of wooden elements on work performance, these studies usually do not investigate the individual effect of VOCs [106]. Studies using wood odors could not show any negative nor positive effects on performances, even if physiological parameters indicated an induction of relaxation [94], [95]. However, one drawback of these studies is that no mechanistical neurophysiological measures as assessed via e.g., EEG could be shown. Future studies could investigate neurocognitive processes related to mental effort that might be increased due to exposure that is perhaps missed on a behavioral level. Further, potential beneficial or even detrimental effects of wood VOCs might also occur on other cognitive levels than working memory. Thus, prospective studies might include a broader range of neurocognitive tests to cover a larger spectrum of potential effect sites.

### 5.5 Conclusion

All in all, the current study does not support the assumption of a positive (e.g., relaxing) nor negative (e.g., irritating) effect of pinewood VOCs in realistic indoor air concentrations. More precisely, participants did not even perceive the exposure at all. While the measurement endpoints used seem to be sensitive to picking up even very slight changes, no concentration-dependent effect could be demonstrated that runs through all results like a common thread. One can, however, conclude that individual factors such as mood or otherwise evoked annoyance might influence ratings which, as has been stressed, are a challenge to handle when intending to regulate olfactory experiences based on subjective or even presumably objective levels (e.g., HR/V). As Greenberg et al., [107] determine, the mere exposure is not a surrogate marker for its perception. Thus, any general regulation of annoying odors/malodors indoors that only relies on VOC concentrations might suffer from the impact of non-sensory factors rendering the intended health protection questionable.

## Supporting information

Supplement to Main Manuscript

## Declaration of Competing interests

All authors declare that they have no personal or financial competing interests.

## Funding Source

The work was supported by funds of the Federal Ministry of Food and Agriculture (BMEL) based on a decision of the Parliament of the Federal Republic of Germany via the Fachagentur Nachwachsende Rohstoffe e. V. (FNR); FKZ: 2220HV026B.

## Acknowledgements

We want to thank Nicola Koschmieder for the organization of and support during the data collection. Further, we thank Lea Drescher, Melina Cosima Mastria, and Andra Nicole Ionescu for their help during the data acquisition. We acknowledge the help of Jörg Reinders, Beate Aust, Michael Porta, Florian Pinnau, and Marion Page for the assistance during the development of the exposure scenarios, the operation of the ExpoLab, and the analysis of the NALF. Lastly, we thank Stefan Kleinbeck for the collaboration on the task development and his help during the data collection.

## Author contributions: CRediT

**Christine Ida Hucke:** Conceptualization, Methodology, Software, Validation, Formal analysis, Investigation, Data Curation, Writing - Original Draft, Writing - Review & Editing, Visualization, Project administration. **Viviane Gallus:** Methodology, Investigation, Writing - Review & Editing. **Katja Butter:** Conceptualization, Writing - Review & Editing. **Julian Elias Reiser:** Methodology, Writing - Review & Editing. **Martin Ohlmeyer:** Conceptualization, Writing - Review & Editing, Funding acquisition. **Christoph van Thriel:** Conceptualization, Methodology, Formal analysis, Writing - Review & Editing, Project administration, Funding acquisition.

## Footnotes

i Note that these approaches might depend on each other and are not proposed to be opposing views but rather connected elements or mechanistic explanations of each other.

## Notes

https://osf.io/kstue/overview?view_only=246c2e54b00d47edaa147bb312451484

### Competing Interest Statement

The authors have declared no competing interest.

