## Supplement to Main Manuscript for "Odor Annoyance, Sensory Irritation or Relaxation: Acute Effects of Real Pinewood Emissions in Indoor Air Scenarios"

BioRxiv Preprint V1 07/26 THIS VERSION OF THE MANUSCRIPT HAS NOT BEEN PEER REVIEWED

### Supplement 1

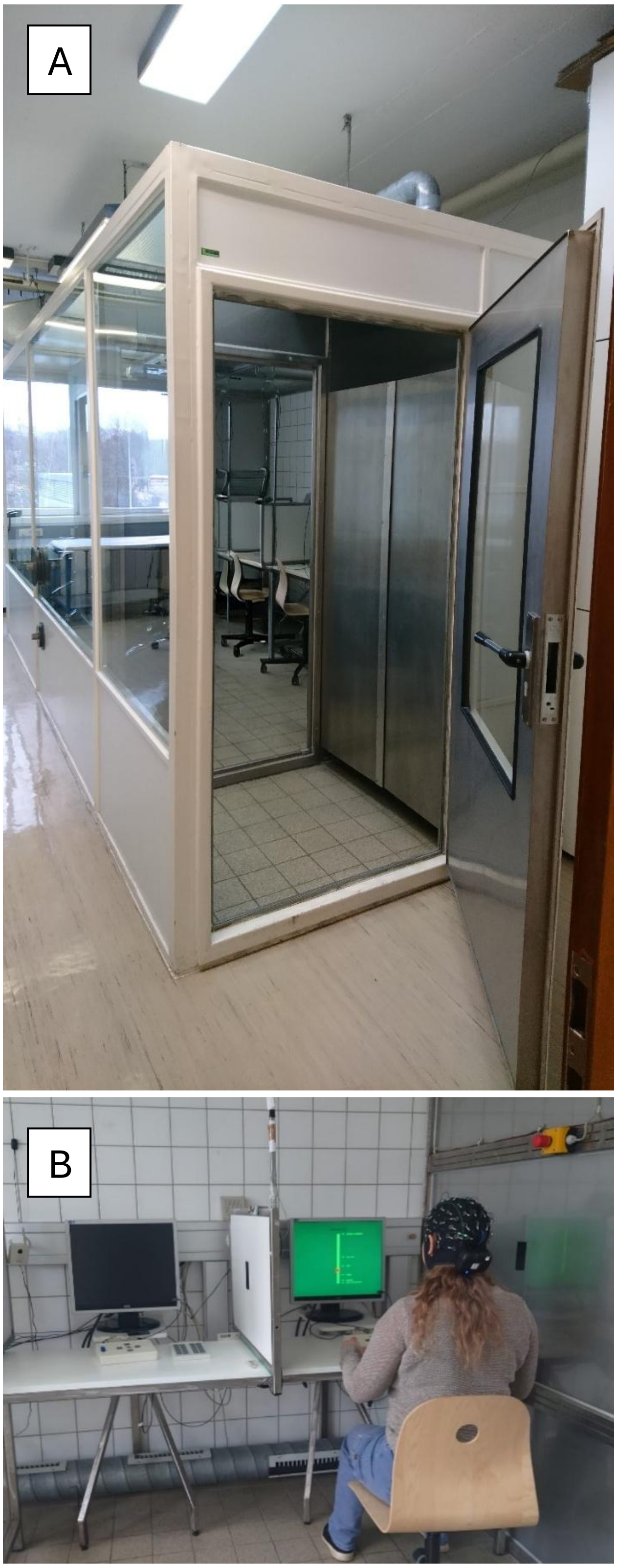


*Figure S 1 (A) ExpoLab with airlock system to prevent changes in indoor VOC concentration. (B) Author sitting in ExpoLab simulating a rating of the indoor air quality on a Labeled Magnitude Scale.*

#### Supplement 2

During the exposure sessions samples of the indoor air were measured using a GC-FID measuring α-pinene and 3-carene in the control room throughout each experiment every 2.5 minutes. Respective average and standard errors are depicted in Figure S 2. Target concentrations of 2 mg/m$^3$ for the GV II and 0.2 mg/m$^3$ for the GV I were closely matched and stable throughout the experiment.

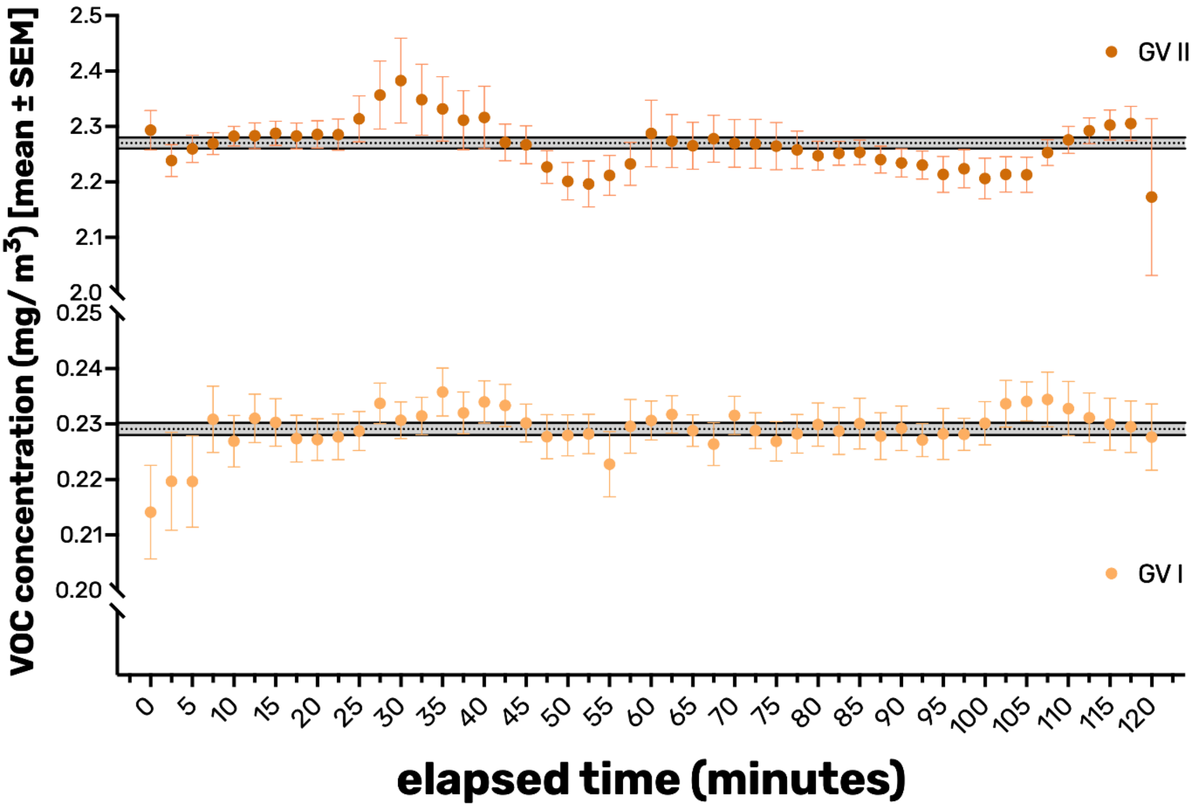


*Figure S 2 Average GC-FID measurements of each exposure session corresponding to the GV I and GV II conditions. VOC: volatile organic compounds, SEM: standard error around the mean. GV: guide value.*

#### Supplement 3A

To establish the exposure protocol and validate the experimental exposures, the concentration and composition of volatile organic compounds (VOCs) in the ExpoLab were additionally determined by active air sampling using Tenax TA® sorbent tubes. For this purpose, a sampling pump operated at a flow rate of 100 mL/min was positioned outside the laboratory. A sampling line was inserted into the exposure chamber through a small opening designed to prevent air exchange between the interior and exterior of the ExpoLab. A Tenax TA® sorbent tube connected to the end of the sampling line was used to collect indoor air samples. Sampling volumes were 500 mL for the GV II condition and 1,000 mL for the control and GV I condition. Following collection, the samples were sent to the Thünen Institute for analysis by thermal desorption gas chromatography–mass spectrometry (TD-GC-MS). Air sampling and analysis of the sorbent tubes were conducted based on DIN ISO 16000-6:2022-03.

As expected, the indoor air was predominantly composed of α-pinene and 3-carene (Figure S 3A). Additional terpenes were detected in minor concentrations, including β-pinene and limonene. Furthermore, benzaldehyde and nonanal were sporadically detected in individual sorbent tubes without a systematic pattern (i.e., occurring across control tubes, GV I and GV II samples). These compounds are unlikely to originate from the indoor air and are likely related to artefacts or degradation products of the sorbent material.

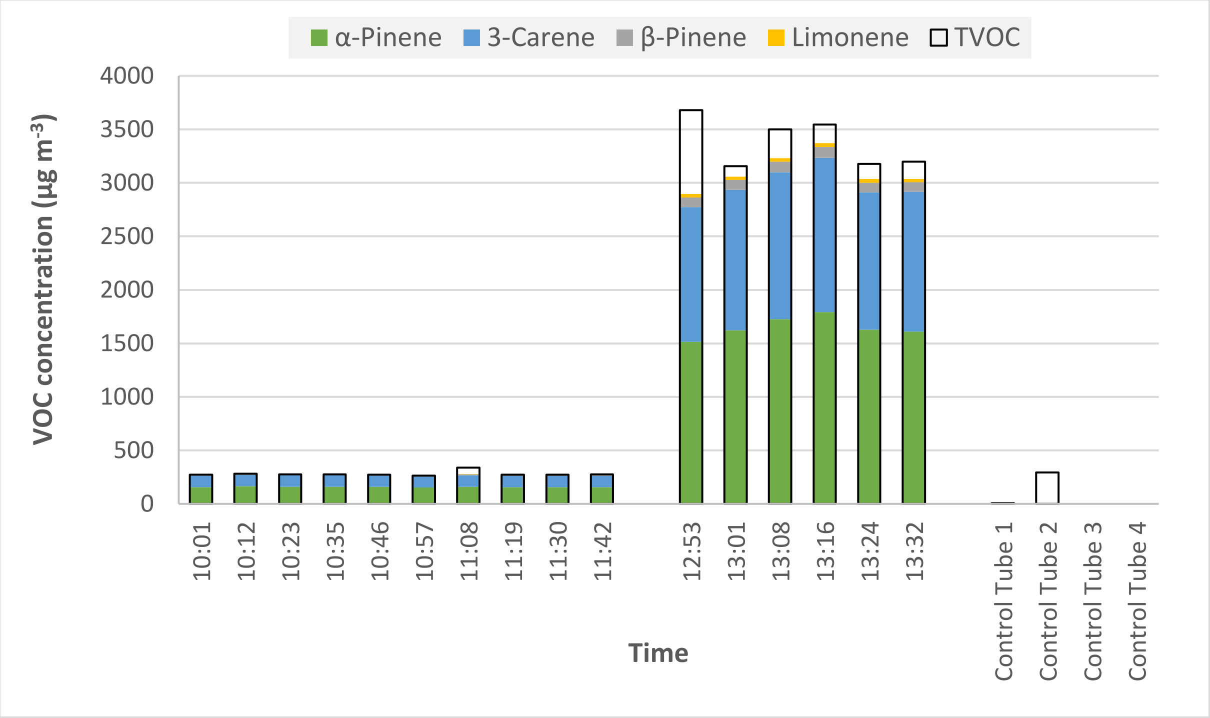


*Figure S 3A Tenax TA®-based VOC measurements during the establishment of the GV I and GV II exposure protocol. Control Tubes: Unopened Tenax TA® sorbent tubes ensuring a plausible and non-contaminated storage and shipping. GV: guide value.*

#### Supplement 3B

During the experiment, each condition (control, GV I and GV II) was sampled twice for VOC composition and concentration following DIN ISO 16000-6:2022. One sample was taken in a morning session and the other one in an afternoon session to control for time of the day. These 6 measurements consisted of three measurement runs each. One at the start of the exposure, one during the break and one at the end, to control for potential variations throughout the exposure session (Figure S 3B).

The VOC composition of the air in the ExpoLab largely corresponded to that observed during the establishment of the experimental conditions (see Figure S 3A), with a slightly higher relative contribution of α-pinene compared to 3-carene.

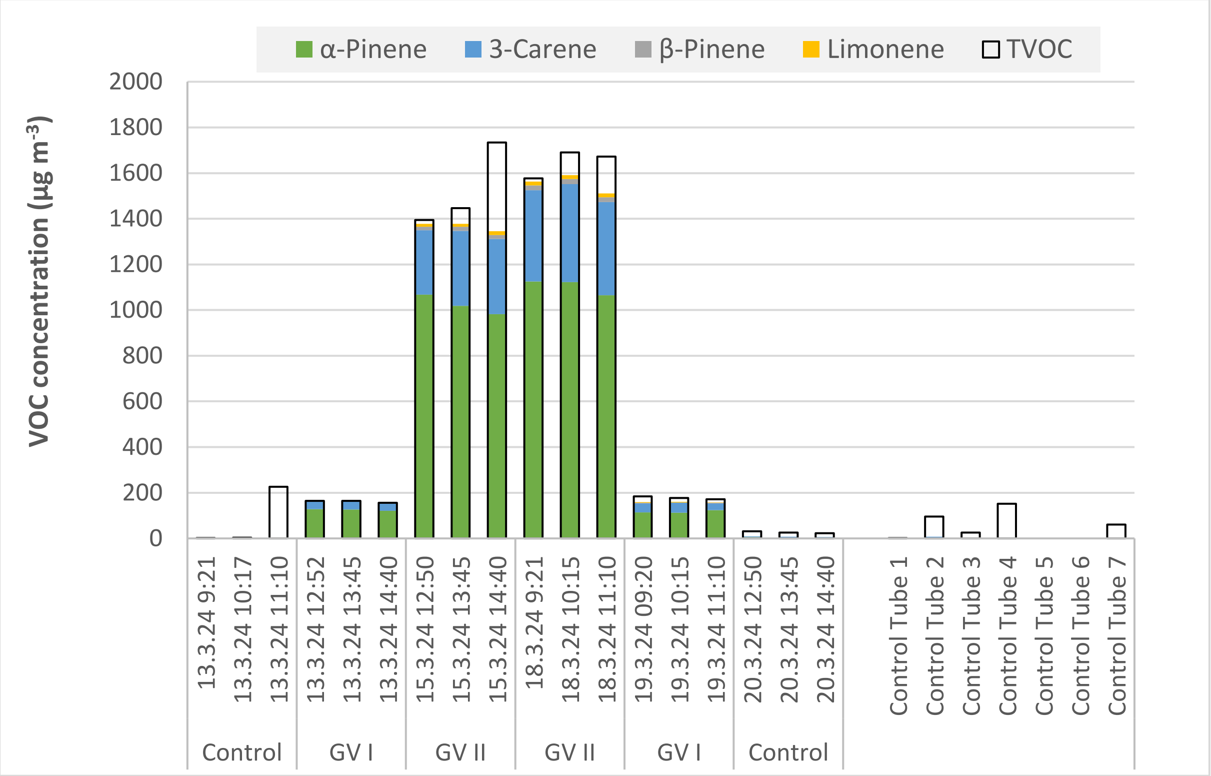


*Figure S 3B* Tenax TA®-based VOC measurements *during the exposure experiment.*
*Control Tubes: Unopened Tenax TA® sorbent tubes ensuring a plausible and non-contaminated storage and shipping.*
*GV: guide value.*

### Supplement 4

Table S 1 indicates mean and standard deviation values of *d'*, corresponding to the 2-back and 3-back tasks.

*Table S 1* Mean (M) and standard deviation (SD) *d'* values for the 2-back and 3-back task

| Exposure | Halves | 2-back | | 3-back | |
|---|---|---|---|---|---|
| | | *d' M* | *SD* | *d' M* | *SD* |
| Control | 1 | 3.48 | 0.76 | 2.34 | 0.63 |
| Control | 2 | 3.43 | 0.60 | 2.41 | 0.72 |
| GV I | 1 | 3.25 | 0.86 | 2.33 | 0.83 |
| GV I | 2 | 3.28 | 0.89 | 2.41 | 0.85 |
| GV II | 1 | 3.32 | 1.00 | 2.33 | 0.90 |
| GV II | 2 | 3.30 | 0.95 | 2.32 | 0.88 |

*Note*. GV = guide value.
